# Structural basis of respiratory complexes adaptation to cold temperatures

**DOI:** 10.1101/2024.01.16.575914

**Authors:** Young-Cheul Shin, Pedro Latorre-Muro, Amina Djurabekova, Oleksii Zdorevskyi, Christopher F. Bennett, Nils Burger, Kangkang Song, Chen Xu, Vivek Sharma, Maofu Liao, Pere Puigserver

**Author notes:** These authors contributed equally: Young-Cheul Shin, Pedro Latorre-Muro. These authors contributed equally: Amina Djurabekova, Oleksii Zdorevskyi. Current address: Department of Chemical Biology, School of Life Sciences, Southern University of Science and Technology; Shenzhen, Guangdong, 518055, China (Young-Cheul Shin). Current address: Institute for Biological Electron Microscopy & Department of Chemical Biology, School of Life Sciences, Southern University of Science and Technology; Shenzhen, Guangdong, 518055, China (Maofu Liao).

## Abstract

In response to cold, mammals activate brown fat for respiratory-dependent thermogenesis reliant on the electron transport chain (1, 2). Yet, the structural basis of respiratory complex adaptation to cold remains elusive. Herein we combined thermoregulatory physiology and cryo-EM to study endogenous respiratory supercomplexes exposed to different temperatures. A cold-induced conformation of CI:III_2_ (termed type 2) was identified with a ∼25° rotation of CIII_2_ around its inter-dimer axis, shortening inter-complex Q exchange space, and exhibiting different catalytic states which favor electron transfer. Large-scale supercomplex simulations in lipid membrane reveal how unique lipid-protein arrangements stabilize type 2 complexes to enhance catalytic activity. Together, our cryo-EM studies, multiscale simulations and biochemical analyses unveil the mechanisms and dynamics of respiratory adaptation at the structural and energetic level.

## Main Text

Brown adipocytes (BAs) are highly specialized cells that respond to lower environmental temperatures by activating adaptive thermogenesis to generate heat and maintain body temperature in mammals(1). Cold-induced sympathetic nervous system activates adaptive thermogenesis in BAs which increases energy expenditure and protects against obesity and diabetes(3). Conversely, BA dysfunction results in defective thermoregulation, obesity and/or metabolic diseases(2, 4–6). Heat production in BAs occurs through two main mitochondrial pathways that depend on Electron Transport Chain (ETC) activity: the classical uncoupling protein 1 (UCP1), which dissipates the proton gradient generated by the respiratory complexes across the intermembrane space (IMS)(5) and a newly identified futile creatine/phosphocreatine cycle, which consumes ATP releasing a molar excess of ADP that increases BA respiration(4). In both cases, ETC activities generate a proton motive force which is utilized to dissipate energy in the form of heat(1, 7). An extensive network of mitochondria with densely packed cristae in BAs provides spatial organization to respiratory complexes and undergoes deep protein and lipid remodeling (2, 8), that in response to low temperatures(2, 9, 10) sustain high oxidation rates of glucose(11), fatty acids (1) and branched-chained amino acids(6). Cold-induced mitochondrial cristae formation and respiration in BAs is regulated by activation of the ER-resident kinase PERK(2, 10). Adipocyte-specific PERK^-/-^ mice are severely cold intolerant and display reduced cristae density and ETC activities, suggesting that architectural cristae and ETC remodeling is indispensable during low temperatures to support mitochondrial respiration(2). However, the structural and dynamic bases of increased mitochondrial respiratory adaptation in response to cold temperatures are largely unknown.

## Cryo-EM revealed conformational transition of respiratory complexes upon cold adaptation

Recent advances in cryo-EM have led to tremendous progress in understanding the structure, assembly, and function of respiratory complexes(12–23). However, previous experimental protocols often require multiple isolation steps, prolonged detergent exposure, and/or utilization of stabilizers to maintain the activity of complexes. For studying catalytic mechanisms, these protocols also commonly add exogenous substrates at saturating concentrations after complex purification. To circumvent these limitations, minimize the impact from detergent-induced delipidation and non-physiological chemicals, and observe endogenous protein complexes bound to relevant substrates, we designed a cryo-EM-based experimental protocol using processed brown adipose tissue samples from mice exposed to different temperature conditions including a genetic model (PERK^-/-^ mice, hereafter KO) (Figure 1). Cold-induced (WT-cold) highly active respiratory complexes and their three-dimensional (3D) structures were determined and compared to those from thermoneutral (WT-TN)(24) and cold acclimated PERK^-/-^ mice (KO-cold)(2) with lower tissue respiratory and thermogenic activity.

**Figure 1.**
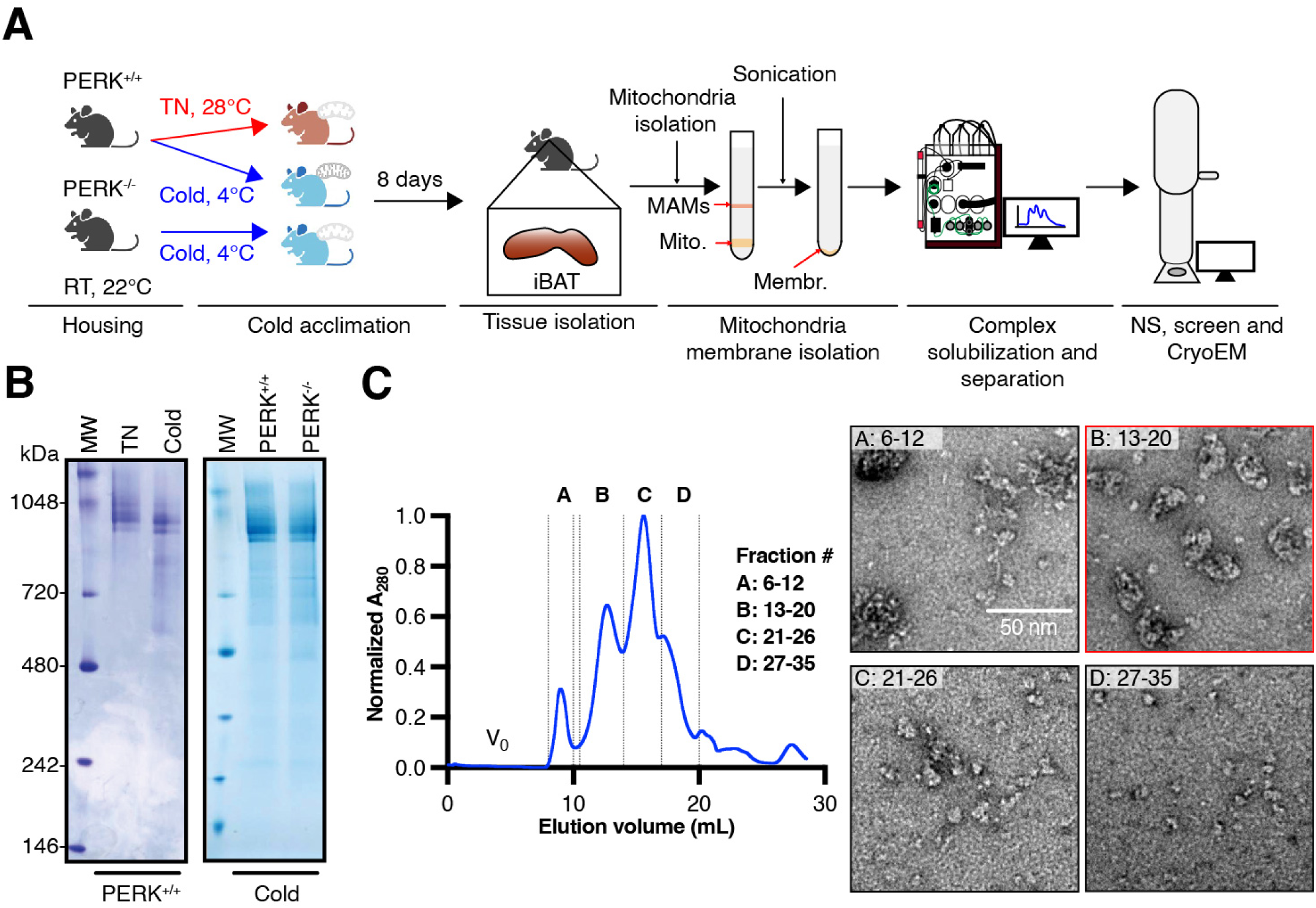
Identification of type 2 CI:CIII_2_ complexes in cold-exposed mouse iBAT. (**A**) Schematic workflow for mitochondrial membrane isolation (see Materials and Methods). Mice were housed at room temperature and exposed to either thermoneutral or cold conditions for 8 days. Interscapular brown fat (iBAT) was isolated, and their mitochondria separated by ultracentrifugation. Mitochondria were sonicated and membranes collected, resuspended in digitonin buffer and complexes separated by size exclusion chromatography before EM analyses. (**B**) Blue-Native PAGE of isolated complexes. (**C**) Size exclusion chromatogram and representative negative stain EM images of each group. Red framed fractions contain respiratory complexes.

Brown fat from mice housed at either 28°C (TN) or 4°C (cold) for 8 days was used to extract respiratory complexes with minimal sample processing (see Materials and Methods). Isolated mitochondria were solubilized with mild detergent digitonin that preserves complex function and structure(25) and immediately fractionated by size exclusion chromatography (SEC) (Figures 1B, 1C and Figure S1) in buffer containing glyco-diosgenin (GDN). The SEC fractions were screened using negative-stain EM, which identified abundant, homogeneous respiratory complexes from the fractions around the second peak in the SEC profiles (Figure 1C, group B). We found few dissociated CI or CIII_2_ across all fractions, indicating predominant assembly of CI and CIII_2_ into supercomplexes. Across all samples, the majority of respiratory complexes were found in fractions 14 to 18 (Figure S1). These fractions were combined, concentrated, and subjected to single-particle cryo-EM. We obtained a series of cryo-EM maps from three samples: WT-cold (types 1A^WT-cold^, 1B^WT-cold^, 2^WT-cold^), WT-TN (type 1^WT-TN^), and KO-cold (type 1^KO-cold^) at the overall resolutions of 3.2-4.1 Å (Figure S2-S4, Tables S1 and S2). Atomic models of complexes in different conformations from these samples were constructed (see Materials and Methods for details). Remarkably, our cryo-EM analysis revealed two major conformations of respiratory supercomplexes that differ in the angle between CI and CIII_2_: one in canonical conformation (termed as type 1) and the other in a non-canonical conformation (termed as type 2 or rotated) (Figure S5 and S6). These two types of complexes were not distinguishable by Blue Native-PAGE (Figure 1B). Type 1 and type 2 conformations were present in the ratio of ∼3:2 in samples from cold acclimated mice and type 2 assemblies were completely absent in thermoneutral and cold-acclimated KO mouse samples (Figure S5), indicating that the type 2 conformation was specifically induced by cold exposure only in WT. Furthermore, cryo-EM analysis of WT-cold sample revealed two 3D classes in type 1 conformation (1A and 1B), one of which demonstrates higher flexibility of peripheral regions of CI and CIII_2_, which would be indicative of the high activity of iBAT upon cold exposure(12, 26). In contrast, WT-TN and KO-cold samples generated only one 3D class in type 1 conformation with more stable peripheral regions. These observations indicate that in response to cold temperatures or conditions of increased energy and thermogenic demands(1, 2), type 1 respiratory complexes transit to structurally distinct type 1 and type 2 conformations.

## Type 2 complexes acquire a planar arrangement associated with lipid remodeling and shortening inter-complex space

Type 2 complexes identified only in cold-acclimated mouse brown fat, show a distinct 25° CIII_2_ rotation around its transversal inter-dimer axis perpendicular to CI membrane domain (MD) (Figure 2 and Figure S7A). Rotation of CIII_2_ results in the opening of the angle between the longitudinal CI membrane domain and CIII_2_ transversal axes as seen from the matrix side (68° vs. 43° in type 1) (Figure 2A). Consequently, one of the CIII_2_ protomers is positioned in closer proximity to CI MD and peripheral arm (PA), while the other is moved further away from their vicinity. On the side and rear views, CIII_2_ accommodates to CI by slight 2° and 9° shifts, respectively (Figures 2C-2D and Movie S1). In order to probe the dynamic behavior of type 1 and type 2 supercomplexes in mitochondrial membrane environments, we constructed large-scale atomic model systems (ca. 2 million atoms) and performed classical atomistic molecular dynamics simulations. In agreement with the cryo EM data, molecular dynamics simulation data show that, relative to CI position, CIII_2_ aligns and maintains a different angular arrangement in type 1 complex compared to type 2 (Figure 2E, top and bottom). Interestingly, the rearrangement of CIII_2_ disrupts all the canonical interactions with CI, but creates new potential contacts between three subunits in CI (NDUFA11, NDUFB7 and NDUFB4) and six in CIII_2_ (Cytochrome C1, UQCRH, UQCR10, UQCRFS1, UQCRQ, UQCRC1) (Figures S7C and S7D) at the matrix and the intermembrane space, with similar topologies as in canonical CI:III_2_ contacts(17, 27, 28). Complementing the structural findings, simulation data reveal that a larger number of protein-protein contacts are maintained in between CI and CIII_2_ in type 1 compared to type 2 complexes (Figure S7B).

**Figure 2.**
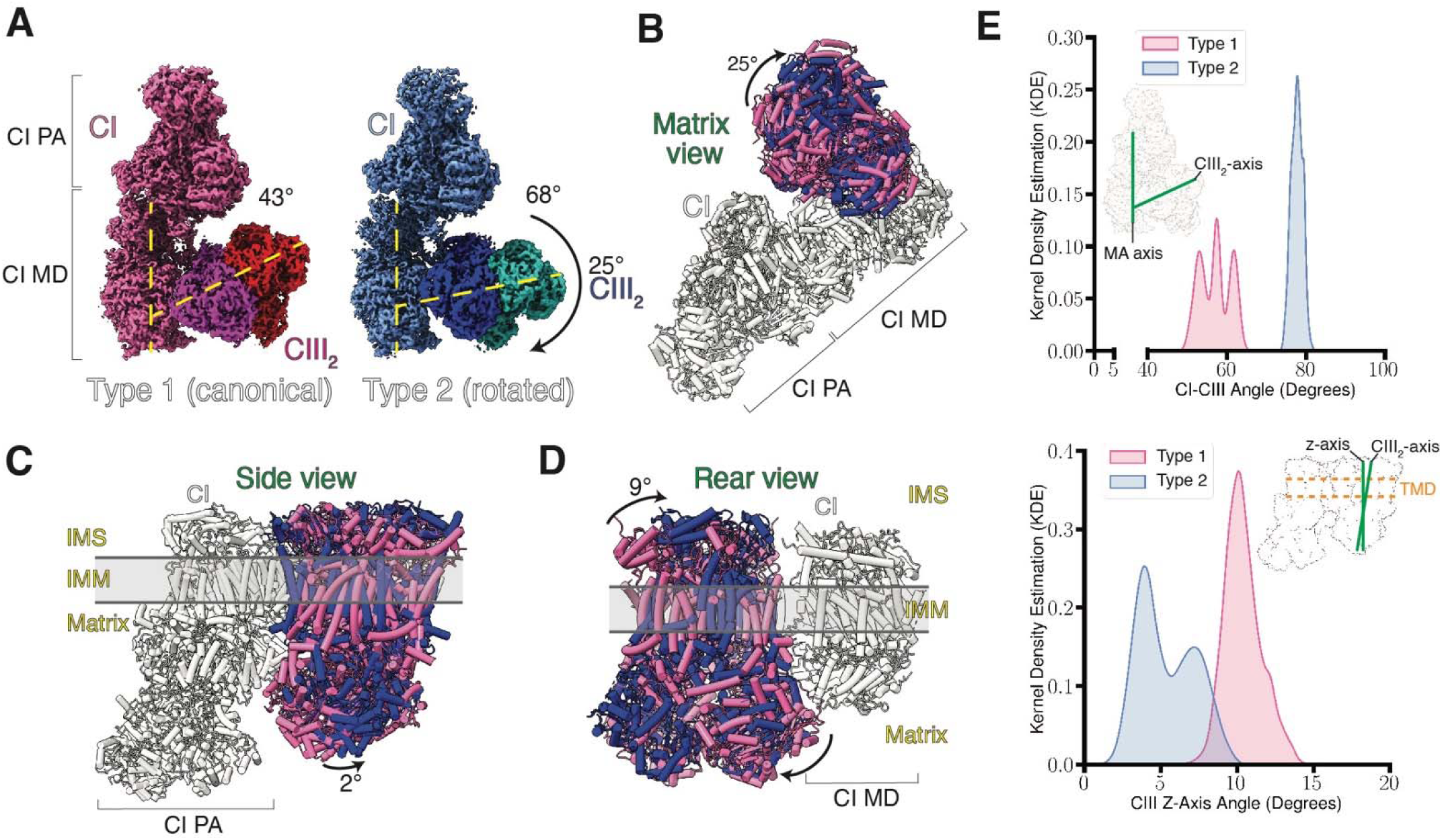
Rotation of CIII_2_ in type 2 complexes upon cold exposure. (**A**) EM density maps (4σ contour) depict the rotation angle of CIII_2_ along CI longitudinal MD axis. Type 1 (hot colors, left) and 2 (cold colors, right) complexes as seen from the mitochondrial matrix side. Each CIII_2_ protomer is colored differently to appreciate contacts with CI. Angles were calculated as the intersection of CIII_2_ transdimer axis with CI membrane domain central axis. (**B-D**), three-dimensional rotation of CIII_2_ in type 2 complexes compared to type 1WT-TN from the matrix (**B**), side (**C**) and rear (**D**) views. Angles are calculated by comparing CIII2 interdimer or cross-sectional axes in type 2 to those in CIII_2_ in type 1. IMM; inner mitochondrial membrane. IMS; intermembrane space between OMM (outer mitochondrial membrane) and IMM. Matrix; matrix compartment of mitochondria. (**E**) Data from 3 x 1 μs molecular dynamics simulations. Top, kernel density estimation (KDE) of the angle between two vectors. The first passes through the membrane arm of CI and the second from the distal to the proximal UQCRC1 subunit of CIII from all simulation replicas of type 1 and type 2. Bottom, kernel density estimation (KDE) of the angle between the z-axis vector with the vector going through the CIII_2_. Data shown is from all three 1 μs simulation replicas of type 1 and type 2 supercomplexes. Green lines indicate the vectors for the calculated angles.

The three-dimensional spatial reorganization of CIII_2_ in type 2 complexes aligns the transmembrane domains (TMDs) of both CI and CIII_2_ (Figure 3A), clearly evident from the absence of tilted CIII_2_. A consequence of TMD alignment is the shortened distances between UQCRB^CIII2^ and NDUFA9^CI^ subunits (∼26 Å vs. ∼12 Å) (Figure 3B), which face the Q exchange compartment (17, 29). Indeed, during molecular dynamics simulations of the type 2 complexes, CIII_2_ tilts ∼ 6° less with respect to membrane normal (Figure 2E, bottom) and a shorter UQCRB^CIII2^-NDUFA9^CI^ distance is maintained (Figure 3C). In addition, CIII_2_ cavities which drive CoQ towards CIII_2_ active site, appear more exposed to CI Q-tunnel exit in type 2 complexes (Figure 3D). We note that a Q molecule indeed has been proposed to bind at the junction of UQCRB^CIII2^ and NDUFA9^CI^ (29). Therefore, in type 2 assemblies, the shortening between UQCRB^CIII2^ and NDUFA9^CI^ may facilitate electron transfer via Q-channeling and/or cycling. This process may be favored by a larger exposure of CIII_2_ active site along with its reduced conformational fluctuations (Figures 2E, top, and 3).

**Figure 3.**
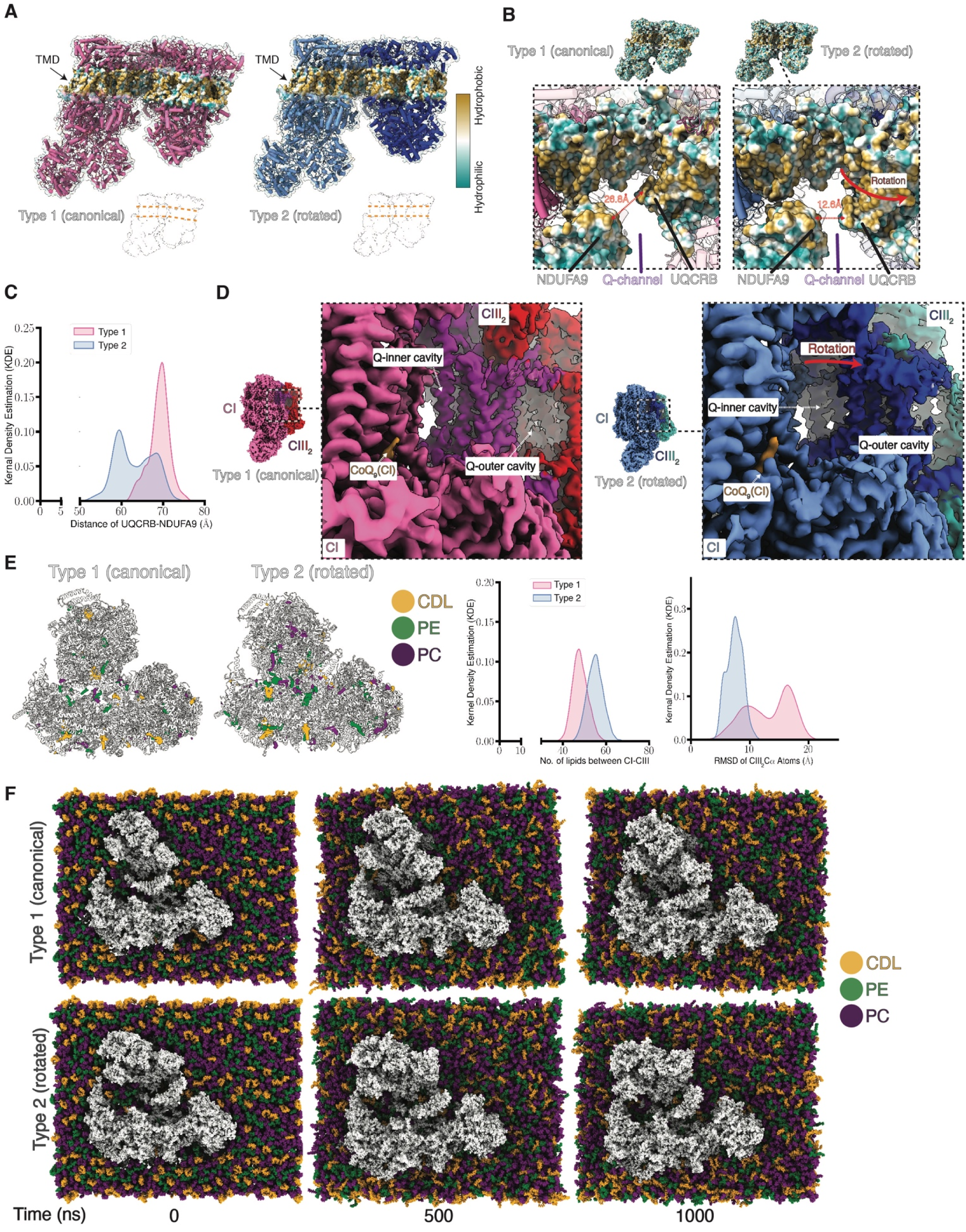
Organization of type 2 CI:CIII_2_ complexes. (**A**) Lipophilicity (golden) across the side view of CI:III_2_ classes shows the transmembrane domain belt that defines straightening of type 2 compared to angled type 1 complexes. (**B**) Rotation of CIII_2_ in type 2 complexes approximates UQCRB (CIII_2_) to NDUFA9 (CI) and forms a hydrophobic protein bridge at the Q-channel. (**C**) Data from 1μs molecular dynamics simulations. The kernel density estimation (KDE) of the distance between center of mass of UQCRB and NDUFA9 for all three simulation replicas of type 1 and type 2 complexes. (**D**) Representation of CIII_2_ CoQ exchange cavities in type 1 (hot colors) and type 2 (cold colors) complexes. Type 2 complexes show complete exposure of both cavities from the CI Q-site view (orange). In B and D, red arrow indicates the displacement distance relative to type 1 complexes. (**E**) Lipid occupancy (isosurface shown at threshold level 0.5 from ChimeraX) in type 1 and type 2 complexes during 3 x 1μs molecular dynamics simulations and depiction of the number of contacting lipids and stabilization of CIII_2_. Left graph, number of lipids at the interface of CI and CIII_2_ (based on the radial distance of 15 Å CI and CIII_2_. Right graph, the kernel density estimation (KDE) of the RMSD of CIII_2_, when the system is aligned based on the membrane arm of complex I for all simulation replicas of type 1 and type 2 complexes. (**F**) Representation of protein-lipid dynamics over 1000 ns.

Cryo-EM along with molecular dynamics simulation data are suggestive of a higher stabilization of type 2 complexes by means that are not fully reliant on protein-protein contacts (Figure S7B). Lipid remodeling in mitochondrial membranes increases ETC activity(25, 30) and sustains cold adaptation and thermogenesis(31). Mitochondrial lipidomics analyses on isolated mitochondria reliably identified 350 unique lipid species (Figure S7E and Table S3). Brown fat mitochondrial membranes from WT-cold adapted mice show a distinct enrichment in unsaturated phosphatidylethanolamine (PE) and phosphatidylcholine (PC) species compared to WT-TN and PERK KO-cold adapted mouse mitochondrial membranes (Figure S7E and Table S3). Molecular dynamics simulations of supercomplexes show that the CI/CIII_2_ interface is more populated with lipids in the rotated conformation of CIII_2_ in type 2 compared to type 1 assemblies (Figure 3E). This arrangement imparts a higher level of dynamics to CIII_2_ in type 1 (Figure 3E), highlighting that a unique conformational state in type 2 is due to distinct lipid-protein arrangements and interactions with flexible lipid PC and PE species(8, 25). In addition, we noticed a trend towards PC species enriching in the vicinity of type 2 complexes (with concomitant decrease in cardiolipin, CDL, population) (Figure S7F) in agreement with the lipidomics data (Figure S7E and Table S3). These results indicate that increasing membrane fluidity through PC and PE lipid species likely provides a platform that allows the assembly and accommodation of type 2 complexes in cold low temperatures(1, 8, 25).

Altogether, our analysis of respiratory complexes from brown fat mitochondria under different thermoregulatory physiological and genetic conditions reveal structural and dynamic changes associated with distinct lipid compositions. Biochemical and *in-silico* evidence suggest that the formation of type 2 complexes with rotated CIII_2_ likely requires the enrichment of flexible lipid species(8, 25) to establish protein-lipid contacts that stabilize and relax CI:CIII_2_ architecture(17) to augment their electron transfer activities(25, 30).

## Enhanced catalytic activity of CI in cold-induced respiratory complexes

Glucose transport in cold acclimated brown fat is nine times higher compared to thermoneutral controls(11). Complete glucose oxidation requires CI oxidoreductase activity and electrons from NADH to reduce ubiquinone into ubiquinol(13). CI peripheral arm (CI PA) open and closed conformations relative to the MD have been extensively reported across species(15, 17, 19, 32) and defined as active and deactive states, respectively. Analyses across our five CI:III_2_ states (type 1^WT-TN^, types 1A^WT-cold^, 1B^WT-cold^, 2^WT-cold^ and type 1^KO-cold^) show distinct global differences representative of each physiological condition. Type 2 complexes display less defined cryo-EM densities at the peripheral regions of CIII_2_, which were observed to a lesser extent in types 1A^WT-cold^ and 1B^WT-cold^ and absent in TN and cold-acclimated PERK KO mice complexes (Figure S7A). Similarly, lower-resolution densities are evident along the PA close to the NADH dehydrogenase domain of CI in types 1A^WT-cold^, 1B^WT-cold^ and especially in type 2^WT-cold^ (Figure S7A) indicative of tilting of CI PA with open/closed states(12). Michaelis-Menten kinetics show ∼2-3-fold increased catalytic efficiency (k_cat_/K_m_) in the WT-cold sample mainly due to increased turnover rates (Figure S8A and B). Moreover, simulations reveal that the PA of type 2 complexes tilts and maintains closer MD-PA angles than type 1 complexes which remain stable in a more open angle conformation (Figure 4A, lower right panel). These results support the increased activity of type 2 assemblies since stable higher opening (larger angle) of CI MD-PA arm may suppress CI’s ability to acquire a closed (smaller angle) state due to the presence of an activation energy barrier(33). On the other hand, narrow swinging of CI PA combined with larger conformational coverage may provide a thermoregulatory advantage in cold temperatures(1, 4, 11, 34).

**Figure 4.**
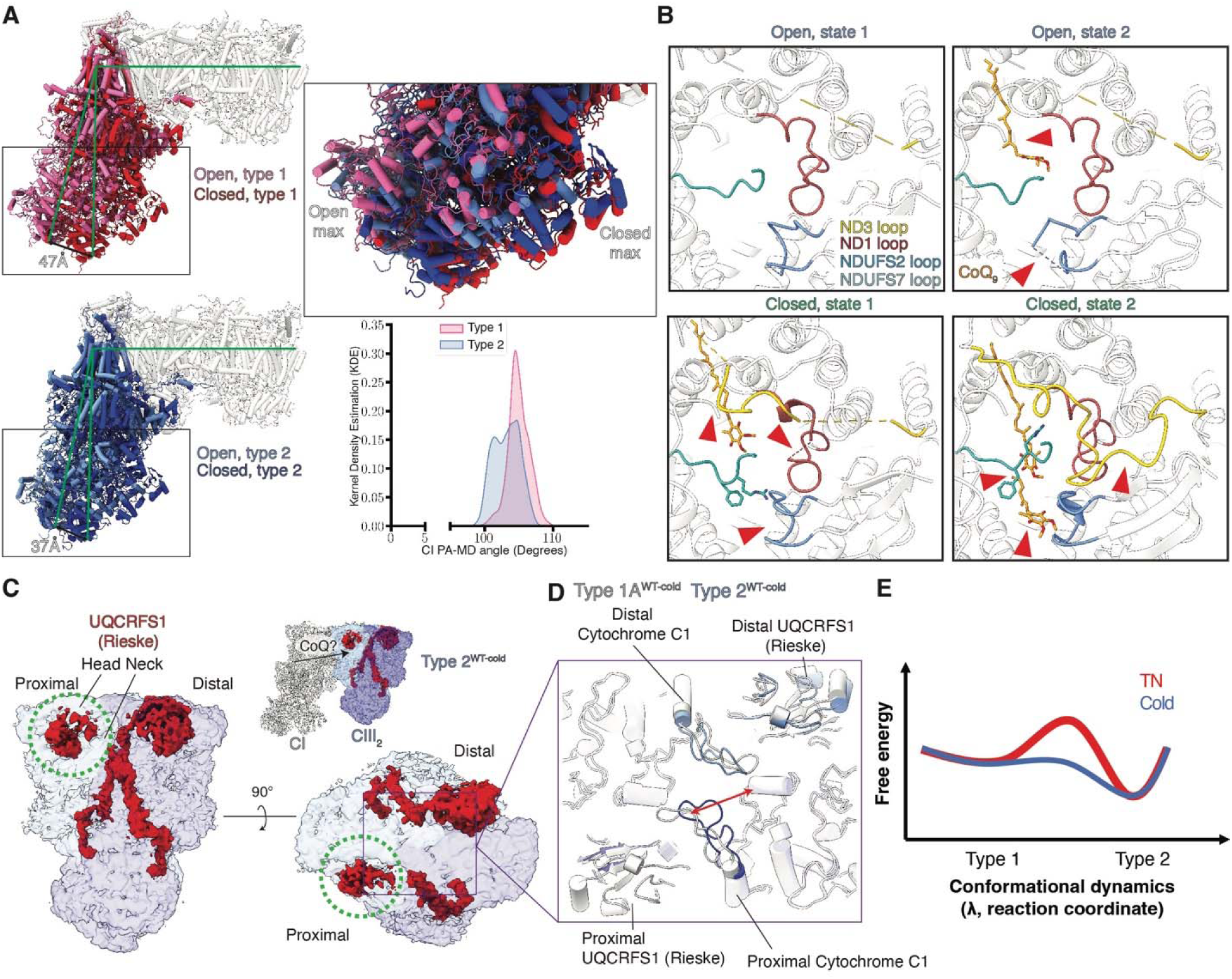
Dynamics of CI and CIII_2_ structures in type 2 assemblies. (**A**) Depiction of maximum opening and closure of CI peripheral arm during 1μs molecular dynamics simulations. Green lines indicate the vectors for the calculated angles. Also, shown is the kernel density estimation (KDE) of the angle between PA and MD of CI from 3 x 1μs simulation data of type 1 and 2 complexes. (**B**) Representation of the active site of CI across different open and closed states in type 2 complexes. (**C**) EM density maps of proximal UQCRFS1 (Rieske) head and neck (green circle) from type 2 complexes show decreased EM densities (red) as seen from the side and IMS views of CIII_2_ (3σ contour). (**D**)Type 2 complexes display increased flexibility of the cytochrome C1 subunit acidic loop (150–170) close to proximal UQCRFS1 subunit and approximates to the same loop in the adjacent protomer. (**E**) Schematic representation of conformational dynamics in TN and cold conditions. An energy barrier in TN-like conditions is minimized in cold-like conditions. As a result of the loss of barrier (blue profile), the type 2 conformation is observed, otherwise not. The low energy type 2 conformation enhances the conformational variation in cold compared to TN conditions.

Focused 3D classification of CI PA corroborates the presence of open and closed states(19) (Figures S8C,D, S9, S10 and Table S4). The majority of particles (63-80% range) show CI PA in a closed conformation. Importantly, type1^WT-TN^ and type 1^KO-cold^ samples, with lower enzymatic activities (Figures S8A and B), showed similar open (20%) and closed (80%) distributions (Figures S8C and S8D). However, in cold-acclimated samples from WT mice, with increased kinetic activity (Figure S8A and S8B), type 1 complexes contained 70% of CI in a closed state and this proportion was even lower in type 2 complexes (63%) (Figure S8C and S8D). Further classifications allowed the determination of two distinct closed conformations and at least one open conformation (Figure S8C and S8D). The closed conformations are predominantly characterized by the ordering of loops and the presence of elongated densities consistent with endogenous CoQ_9_ at the active site of CI (Figure S11 and S12). CoQ_9_ occupies Q-tunnel of CI in type 1B^WT-cold^ and type 2 assemblies and to a lesser extent in type 1^KO-cold^ samples (Figure S11A) with the benzoquinone ring positioned at the Q deep (Q_d_) chamber and the isoprenoid tail occupying the tunnel, consistent with previous reports(15, 20, 32). Remarkably, two different open conformations were observed in type 2 assemblies which illustrate the entry and accommodation of CoQ_9_ during catalysis in a stepwise manner (Figure 4B, state 1 transitions to state 2, top panels). In the open state 2, CoQ_9_ penetrates the Q tunnel from the membrane milieu, rests at the shallow (Q_s_) chamber and triggers the ordering of the NDUFS2 loop at the Q_d_ chamber (Figure 4B, top right). Then, CI transitions to a closed state which fully organizes the NDUFS2 loop, promotes the flipping of the ND1 loop and partially organizes the ND3 loop (Figure 4B, bottom left). Finally, CoQ_9_ moves to the deep chamber (Q_d_) upon flipping of the NDUFS8 loop, and the ND3 loop is fully ordered reaching a fully competent catalytic state (Figure 4B, bottom right). Our findings provide detailed mechanistic and dynamic insights into the intermediate states of the catalytic cycle of CI (19–21, 23, 32, 33, 35).

Important structural differences are seen in the binding modes of CoQ_9_ head group in type 2^WT-cold^ and type 1^WT-TN^ consensus maps. In the latter, the head group is within electron transfer distance (∼11 Å) of terminal FeS cluster N2 (yielding a fast time constant of ca. 200 ns). Interestingly, the rearrangement of loops occupying the Q tunnel brings the sidechain of conserved Arg122 of NDUFS8 loop in hydrogen bonding distance to the Q head group. Previous structural and biochemical data have highlighted the importance of arginine in complex I activity(36). Hybrid QM/MM (quantum mechanical/molecular mechanical) molecular dynamics simulations show rapid proton transfer(s) from Tyr141 (NDUFS2) and/or Arg122 upon two electron reduction, but not on one electron transfer to Q, forming (anionic) quinol, commensurate with the fast rate of electron transfer between N2 and CoQ_9_ (Figure S8E, Tables S5-S7). On the other hand, the CoQ_9_ head group in catalytically more active type 2^WT-cold^ state is located ca. 15 Å from the N2 FeS cluster, which will drop the electron transfer rate to hundreds of microseconds (Figure S8E, Tables S5-S7). The latter site most likely prevents the reverse electron transfer(37–39), and thus enhances the rate of forward reaction of complex I. Alternatively, it has been suggested based on structural data that more than one Q molecules can occupy the Q tunnel(32) and complex I may function with the involvement of two Q molecules with a redox reaction between them(40). To further probe the electron distribution for each complex type in this alternative experimental setting, we created additional QM/MM setups with two quinone molecules modeled in the Q tunnel (see Materials and Methods). We modeled a CoQ_1_ at site 1, and another CoQ_9_ in the site 2/3 region of the Q tunnel(32). Based on type 2^WT-cold^ consensus, CoQ_1_ was modeled at site 1^F^, and a CoQ_9_ in site 2/3. After adding the first electron to the system, in type 1^WT-TN^ the spin density almost fully concentrates on CoQ_1_ at site 1 (Figure S8F, top left panel), while remarkably in type 2^WT-Cold^, site 3 (occupied by CoQ_9_) becomes a preferential location for the electron (Figure S8F, bottom left panel). This suggests that in type 2^WT-Cold^ conformation, exchangeable CoQ_9_ can accept electron from Q species bound at site 1^F^. Upon modeling the two-electron reduction step (Figure S8G), we find that a spontaneous proton transfer occurs from Arg274^ND1^ to CoQ_9_ located at site 3, while CoQ_1_ at site 1^F^ does not accept protons. Also, to note is that the electron transfer rate between the N2 cluster and CoQ molecule at site 1 is orders of magnitude higher than when CoQ is at site 1^F^ (Table S7). Interestingly, type 2^WT-Cold^ arrangement of Q molecules leads to an exceedingly high rate of electron transfer for the CoQ_1_ – CoQ_9_ pathway (Table S7). All in all, our QM/MM simulations based on cryo EM data suggest how type 2 assemblies may achieve higher rates of catalysis in two different CoQ settings in the Q tunnel of CI, in agreement with the kinetic data from cold-stimulated samples.

Even though the open and closed states of CI have been defined as deactive and active states, respectively, current research has proposed that the open-to-closed conformations are part of CI catalytic cycle(12, 33). Our experimental and *in-silico* results indicate that higher kinetic activities correlate with more even proportions of open and closed conformations (Figure S8C and D) which would be indicative of more frequent transitions(12) as biochemically observed(41). In addition, our findings support a role for the open/closed transition during the catalytic cycle of CI as an energetically modulable state(1, 12, 21, 33) that defines the efficiency of electron transfer (Figures S8E-S8G). This efficiency may be favored in type 2 complexes by the narrower opening angles of CI arm (Figure 4A) that minimizes the energy loss during each cycle to enhance ETC-dependent nutrient oxidation and augment heat production in response to cold environments(1, 7).

## Two CIII monomers in type 2 assemblies display structural and catalytic variations

Respiratory CIII_2_ is a homodimer that oxidizes CoQH_2_, which can be provided by CI, and transfers these electrons to cytochrome C through the UQCRFS1 (Rieske)-Cytochrome C1 subunits(13). Each dimer contains two oxidative sites (Q_p_) and two reductive sites (Q_n_) connected through a network of six heme molecules that are acceptors and donors of electrons. However, under physiological conditions, the structural mechanisms of electron transfer from CoQH_2_ to cytochrome C and how each CIII monomer contributes to the catalytic cycle of CoQ oxido-reduction remains unclear (13, 42). We find that despite their rotation relative to CI, CIII_2_ complexes in type 2 respiratory assemblies show a similar overall structure compared to CIII_2_ in type 1, or canonical, assemblies. Identical heme distribution across types 1 and 2 indicate preserved symmetry of the dimer across states (Figure S13A). However, type 2 complexes have weaker densities at the head and neck of UQCRFS1 proximal to CI, suggestive of higher levels of mobility, activity, and transfer of electrons from the reduced CoQH_2_ to Rieske (Iron-sulfur cluster) and heme c in cytochrome C1 subunit (Figure 4C and Figure S13B)(26). In contrast, distal UQCRFS1^CIII2^ head and neck domains seem stabilized by NDUFB7^CI^ (Figure S7C).

Adjacent to mobile UQCRFS1^CIII2^, type 2^WT-cold^ complexes show swinging of the acidic loop in cytochrome C1 subunit (^150^EEVEVQDGPNDDGEMFMRPGK^170^) approximating to its sibling cytochrome C1 loop from the other CIII (Figure 4D and Figure S13C). Conversely, fluctuations of cytochrome C1 loop in other states from the cold condition are minor (types 1A^WT-cold^ and 1B^WT-cold^) or missing in complexes from low respiratory conditions (types I^WT-TN^ and I^KO-cold^) (Figure 4D and Figure S13C). Given the acidic nature of these loops, relocation of these towards each other would be possible through protonation of aspartates and glutamates as a consequence of higher ETC activity during cold acclimation^5^.

Each CIII monomer contains two CoQ sites: one is oxidative (Q_p_) and the other is reductive (Q_n_). It is unclear whether each CIII monomer can perform simultaneous oxido-reductase activities or be only either oxidative or reductive(13). At CIII_2_ active sites, identifiable CoQ densities are distributed at the Q_p_ (oxidative) and Q_n_ (reductive) sites (Figure S13D). A total of 4 molecules are observed in all states except for type 2 complexes, where CoQ is absent from the distal oxidative (Q_p_) site (Figure S1D). At this location, distal UQCRFS1 is stabilized by NDUFB7 (Figure S7C), demonstrating more defined densities than the proximal subunit in type 2 assemblies (Figure 4C). The observations of higher flexibility of UQCRFS1 and the swinging of cytochrome C1 loop indicate higher oxidative activity of the CIII protomer proximal to CI. This denotes specialization of each CIII protomer where proximal CIII is oxidative and distal is reductive, consistent with previous studies(13, 17, 42).

## Discussion

The structural mechanisms that allow respiratory complex adaptation to increased energetic demands remain largely unexplored(43). In response to cold temperatures, brown adipose tissue initiates adaptive thermogenic responses that are reliant on higher respiratory function to sustain high nutrient oxidation rates(1, 2, 7, 44). Consistently, cold exposure in WT mice increases respiratory complex activity (Figure 4A and Figure S7A, S8A and S8B) but more importantly, induces the formation of type 2 or non-canonical CI:III_2_ complexes (Figure S7A)(1, 2, 7). CI:III_2_ complexes with rotated CIII_2_ have been observed, but the low resolution of the maps and absence of physiological and genetic context precluded the analysis of their functional significance(26, 29, 45). Survival to cold temperatures requires remodeling of brown fat mitochondrial protein and lipid composition to increase cristae density and maximize ETC activities(2, 31). In this process, type 2 complexes may form during exposure to cold temperatures after protein-protein interactions are released allowing new contacts to stabilize the type 2 conformation to fit cristae architecture (Figure 3A)(2, 44) therefore explaining type 2 complex enrichment in brown fat relative to other assemblies in resting state tissues (26, 29, 45). Rotation of CIII_2_ in type 2 complexes results in the planar arrangement of CI:III_2_ TMDs which correlates with the straightening of mitochondrial cristae during cold acclimation(44), perhaps explaining the high abundance of type 2 CI:III_2_ assemblies in active BA mitochondria (2, 44)(Figures 3A and 3B). Similar correlation between cristae morphology and respiratory complex shape was recently reported in *T. thermophila*(42) and *A. thaliana*(28), without CIII_2_ rotation, suggesting a general adaptive mechanism across species(25). Type 2 complexes may localize at specific subregions of mitochondrial cristae that undergo elongation and planar arrangement during cold temperatures(2, 44), a process that cooperatively may benefit from the presence of flexible lipid species such as PC and PE that stabilize respiratory complexes (Figures 3E, 3F, S7E and S7F)(8, 25). Therefore, we suggest a potential mechanism of transition from type 1 to type 2 under cold conditions that occurs by lipid remodeling (PC/PE) that increases the flexibility of the membranes(25) and the formation of lipid-protein rearrangements that allow the accommodation of type 2 complexes to straightened cristae, resulting in lowering of energy barrier when compared to thermoneutral conditions (Figure 4E).

CI and CIII_2_ exert a repressive effect on each other limiting their activity(17). Molecular dynamics simulations of full supercomplex in realistic membrane environment reveal type 1 conformation is dynamic, but uniquely different from type 2 which shows clear stabilization of CI and CIII_2_ independently (Figures 3B, C and E). The architecture of type 2 complexes may be suggestive of more relaxed (energetically favorable) CI:CIII_2_ assembly that reduces those repressive forces to promote activity (Figures 2 and 4E). This view is supported by i) the tilting of the peripheral arm (Figure 4A)(12) and landscape of conformations at the active site of CI (Figure 4B), and ii) the increased flexibility of the UQCRFS1 subunit proximal to CI (Figure 4D)(26) along with swinging of its adjacent loop in cytochrome C1 (Figures 4D and E) in CIII_2_.

Mechanistically, the narrower opening of CI peripheral arm would sustain an energetically favorable momentum that facilitates frequent tilting events and higher catalytic efficiency (Figures 4A and 4E). Additionally, the reduction of inter-complex distances from CI to CIII_2_ at the UQCRB-NDUFA9 protein bridge in type 2 complexes (Figures 3B and 3C) likely participate in efficient electron transfer via Q-channeling (17), as opposed to CoQ membrane diffusion(29). We propose that the formation of type 2 assemblies with rotated CIII_2_ increases CI activity and proton pumping to the IMS. Increased proton presence subsequently protonates cytochrome CI^CIII2^ acidic loops at the IMS and favors their proximity, which augments proximal UQCRFS1^CIII2^ (Rieske) flexibility and its ability to transfer electrons to cytochrome C(13). Remarkably, the asymmetries observed in CIII_2_^type^ ^2^ (UQCRFS1 head and neck, cytochrome C1 loop, and CoQ distribution) (Figures S13B-D) indicates that each protomer performs either CoQ oxidative or reductive activities, a conserved mechanism across species to maximize complex activity(26, 42). Therefore, type 2 complex stoichiometries are modulated under cold temperatures to enhance ETC function and respiratory adaptation in BAs.

Our structural data indicate that upon rotation of CIII_2_, type 2 assemblies are highly active in the context of cold adaptation to increase respiration that supports heat production and body temperature. This model is also supported by previous structural and computational studies(12, 15, 17, 20, 21, 23, 32, 42, 46) and reveals structural and catalytic insights in our understanding of CI:III_2_ mechanism and dynamics in adaptation to energetically demanding cold temperatures.

---

**Figure S1.**
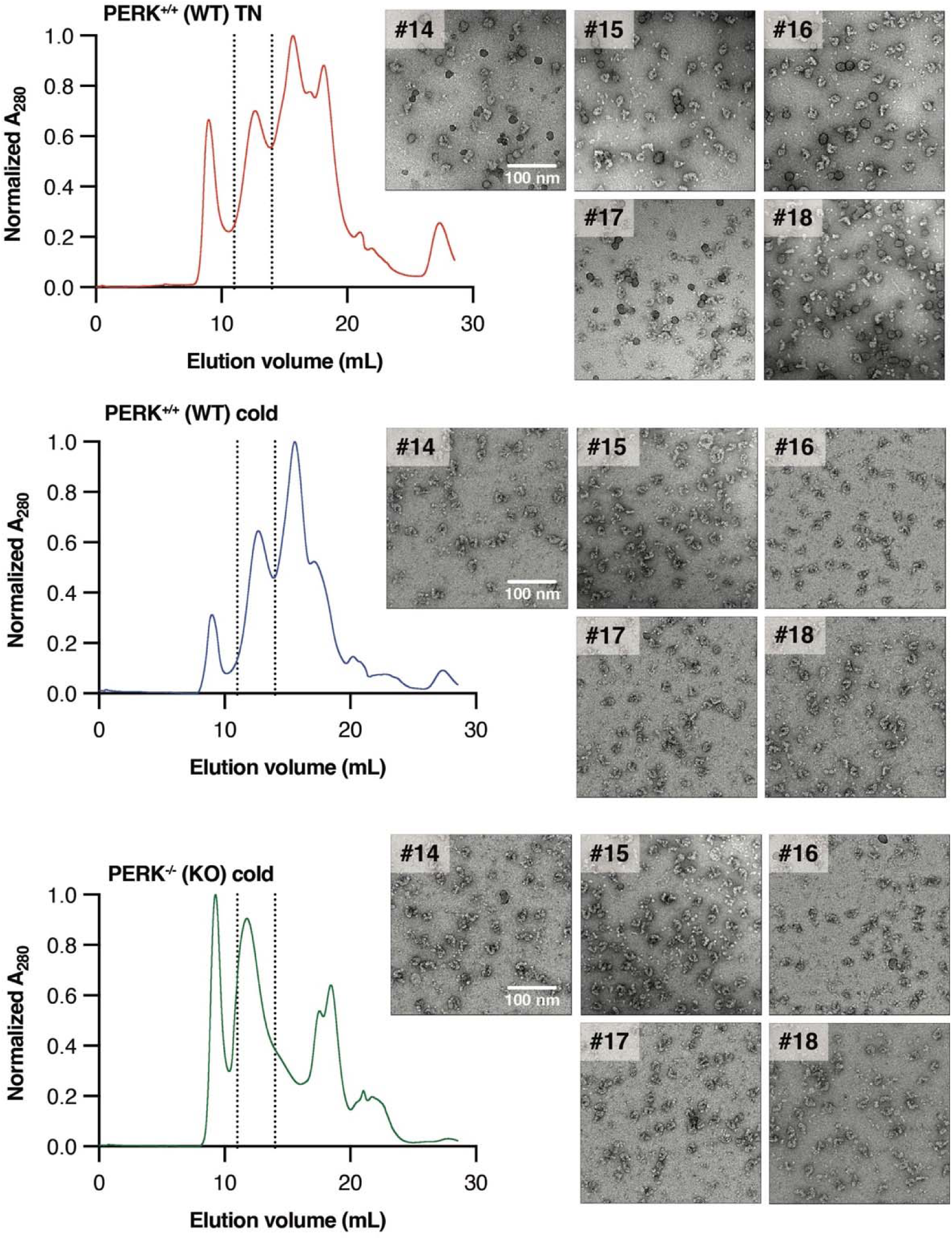
Screening of negative-stain EM (NS-EM) fractions for cryo-EM analysis. Digitonin-solubilized proteins were resolved by size exclusion chromatography in a Superose 6 column and fractions were subject to NS-EM analysis. Indicated fractions were pooled and used for Cryo-EM analysis. Complexes from different conditions were isolated in parallel. PERK^+/+^; wild type mice. PERK^-/-^; adipocyte-specific PERK KO mice.

**Figure S2.**
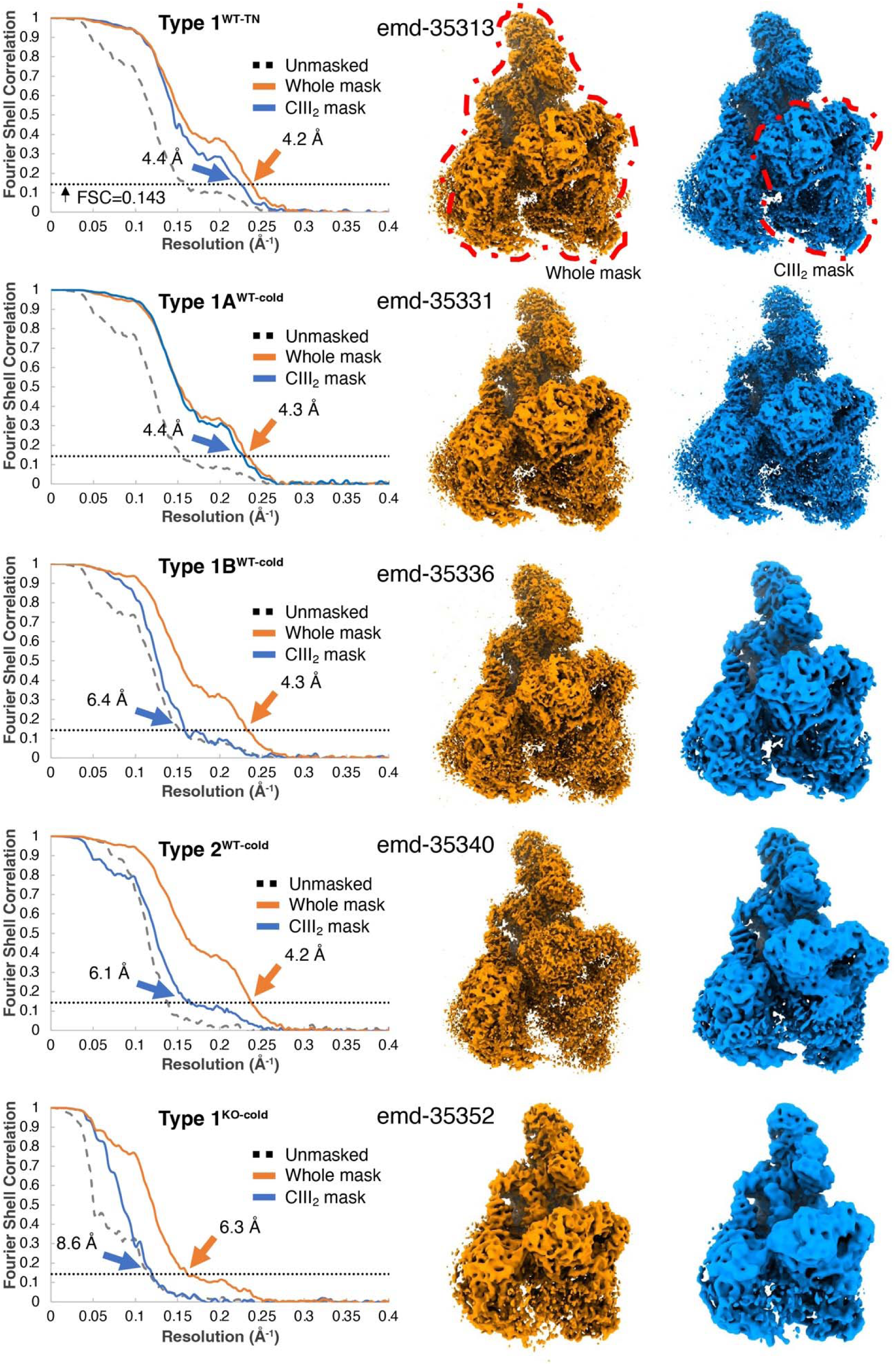
Fourier shell correlation (FSC) curves and full maps postprocessed using full mask and CIII_2_ mask. Unmasked (dashed black line), masked with full mask (solid orange line) or masked with CIII_2_ mask (solid blue line) gold standard half-map FSC are shown with FSC cutoff 0.143. Maps were refined using “relion_postprocess”. These maps were utilized for specifying the location of CI (orange) or CIII_2_ (blue) after focused refinement on peripheral arm, membrane domain or CIII_2_. Maps provided in EMDB are full maps with full mask (‘full map’ in Table S2).

**Figure S3.**
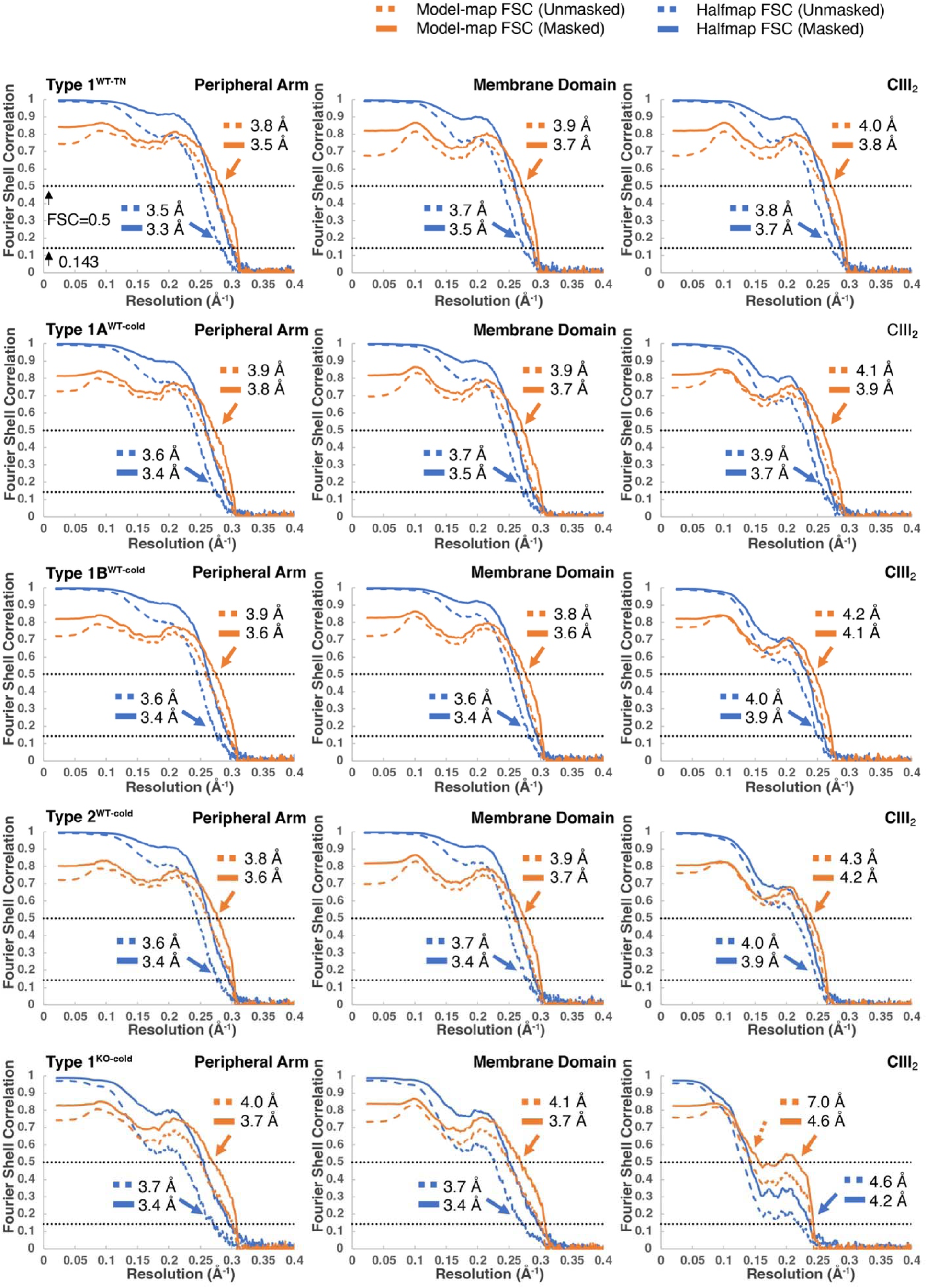
Fourier shell correlation (FSC) curves of focused refined maps. The FSC curves for three focused refined maps with masks (peripheral arm, membrane domain and CIII_2_) in five states. The gold standard half-map FSC and model-map FSC are shown as blue and orange lines, respectively (dashed line for unmasked and solid line for masked). The gold standard FSC cutoff (0.143) and model-map cutoff (0.5) are indicated with dashed black lines and estimated resolution on the cutoffs are indicated with colored arrows.

**Figure S4.**
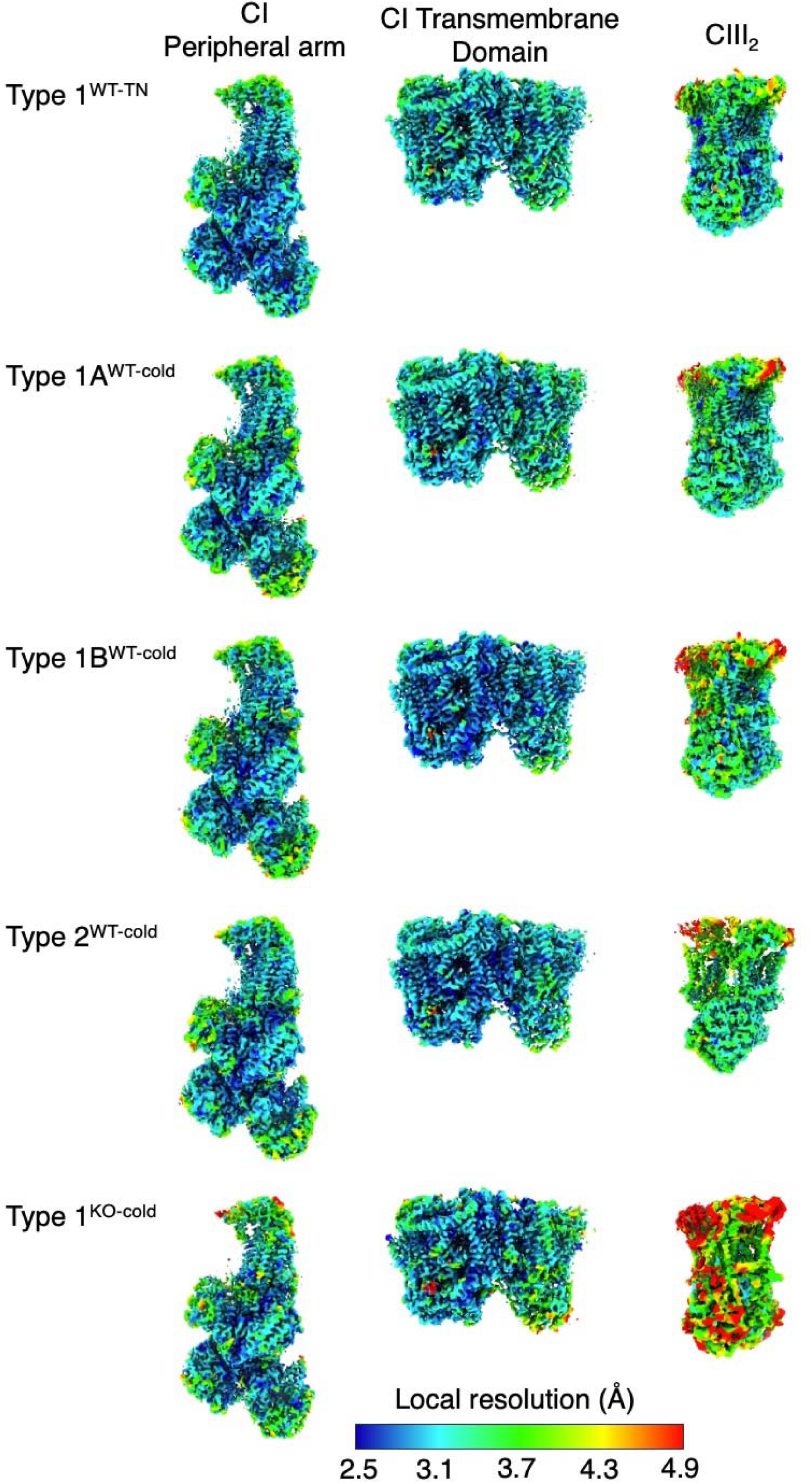
Local resolution estimates of focused maps across states. Determination of local resolution estimates for three focused refined maps with masks (peripheral arm, membrane domain and CIII_2_) in five states.

**Figure S5.**
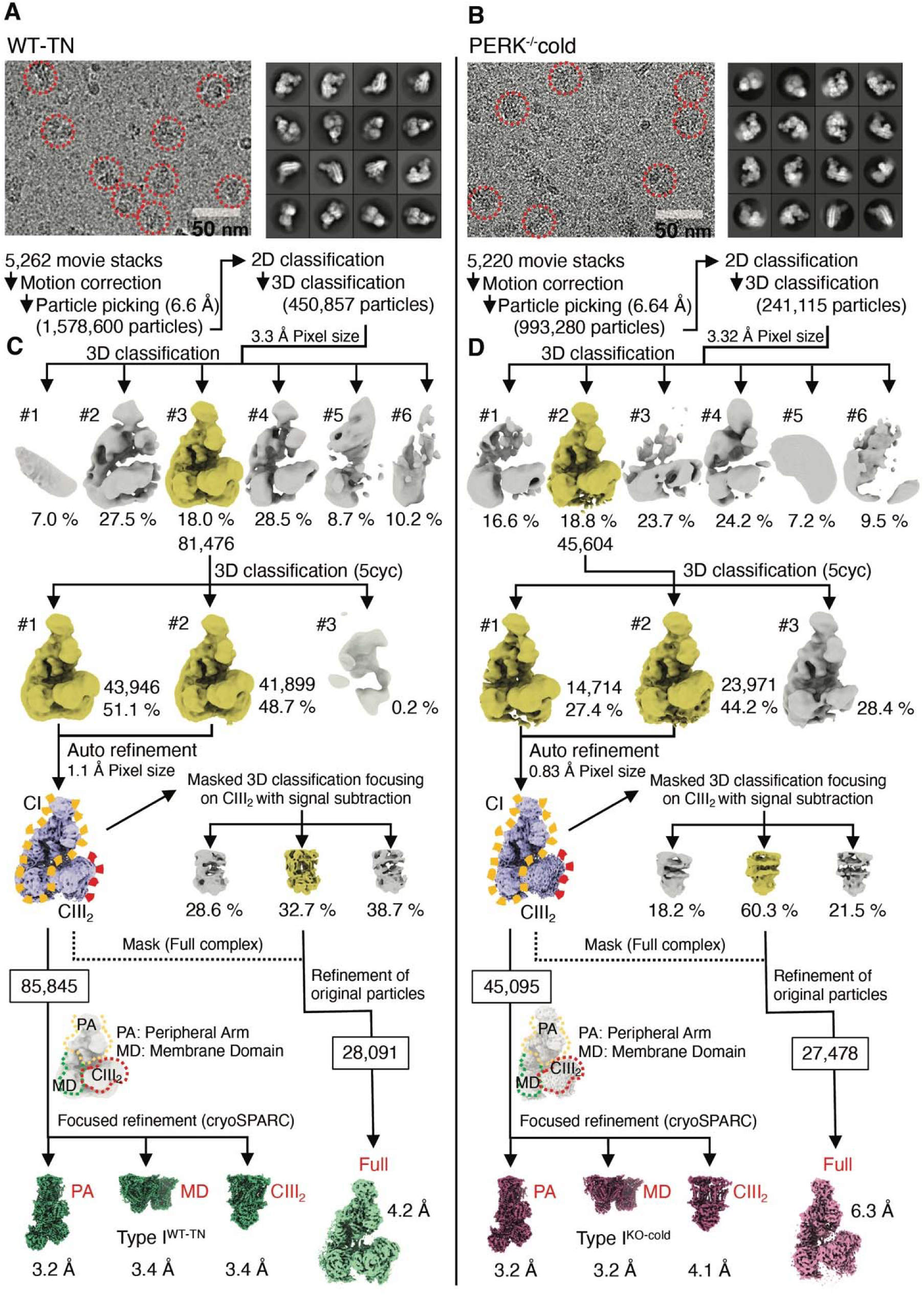
Image processing of respiratory complexes from PERK^+/+^ thermoneutral (TN)-acclimated and PERK^-/-^ cold-acclimated mice. (**A** and **B**) Representative micrograph, 2D classes and process of particle sorting with binned (6 for TN and 8 for PERK-/− cold, respectively) images. The particles red encircled indicate supercomplex. (**C** and **D**) Work-flow of image illustrates the presence of a single 3D class under these conditions (C for TN and D for PERK-/− cold, respectively).

**Figure S6.**
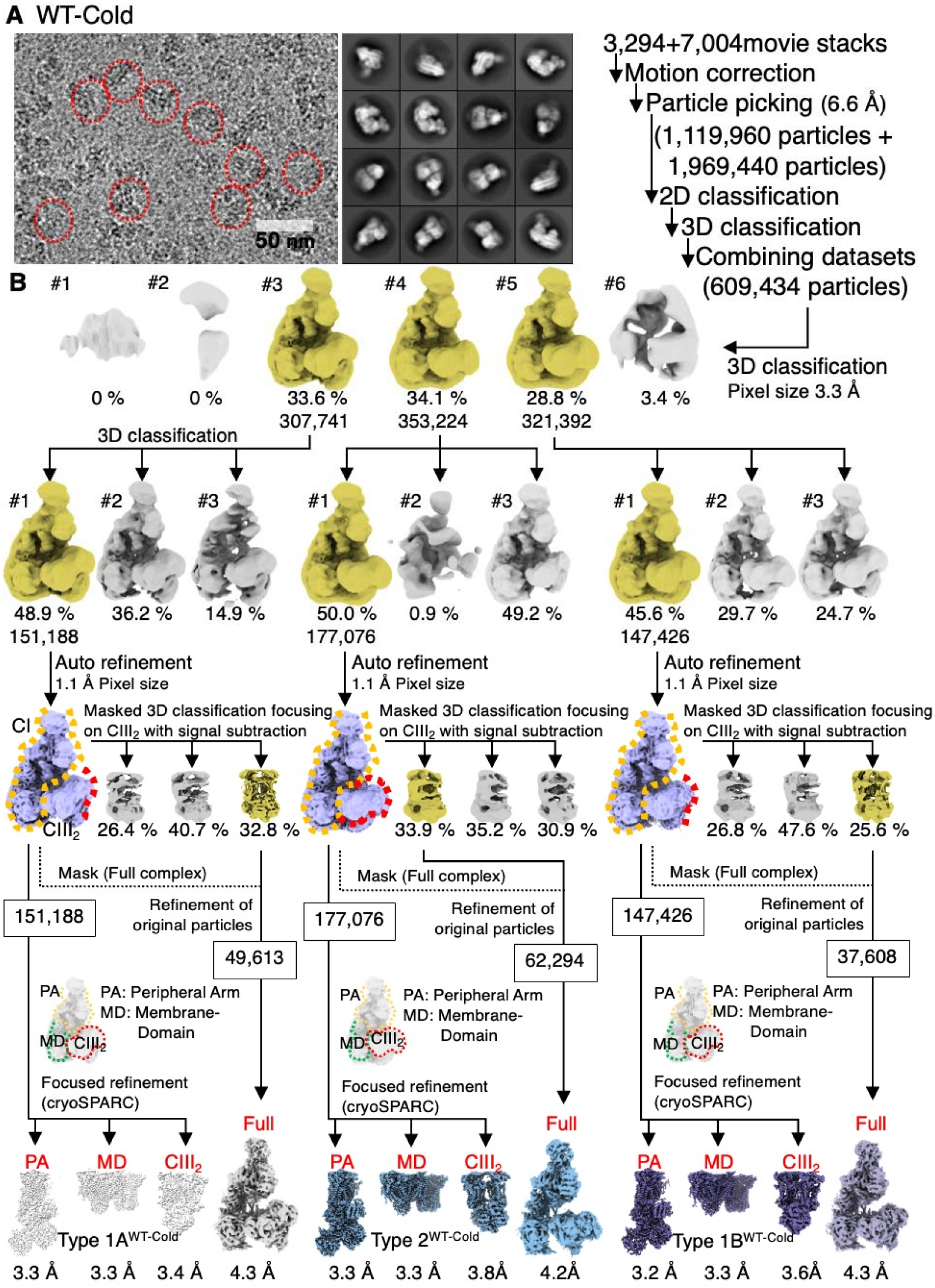
Image processing of respiratory complexes from PERK^+/+^ cold-acclimated mice. (A) Representative micrograph, 2D classes and process of particle sorting with binned 6 images. The particles red encircled indicate supercomplex. (B) Work-flow image analysis showing the presence of multiple 3D classes in this condition.

**Figure S7.**
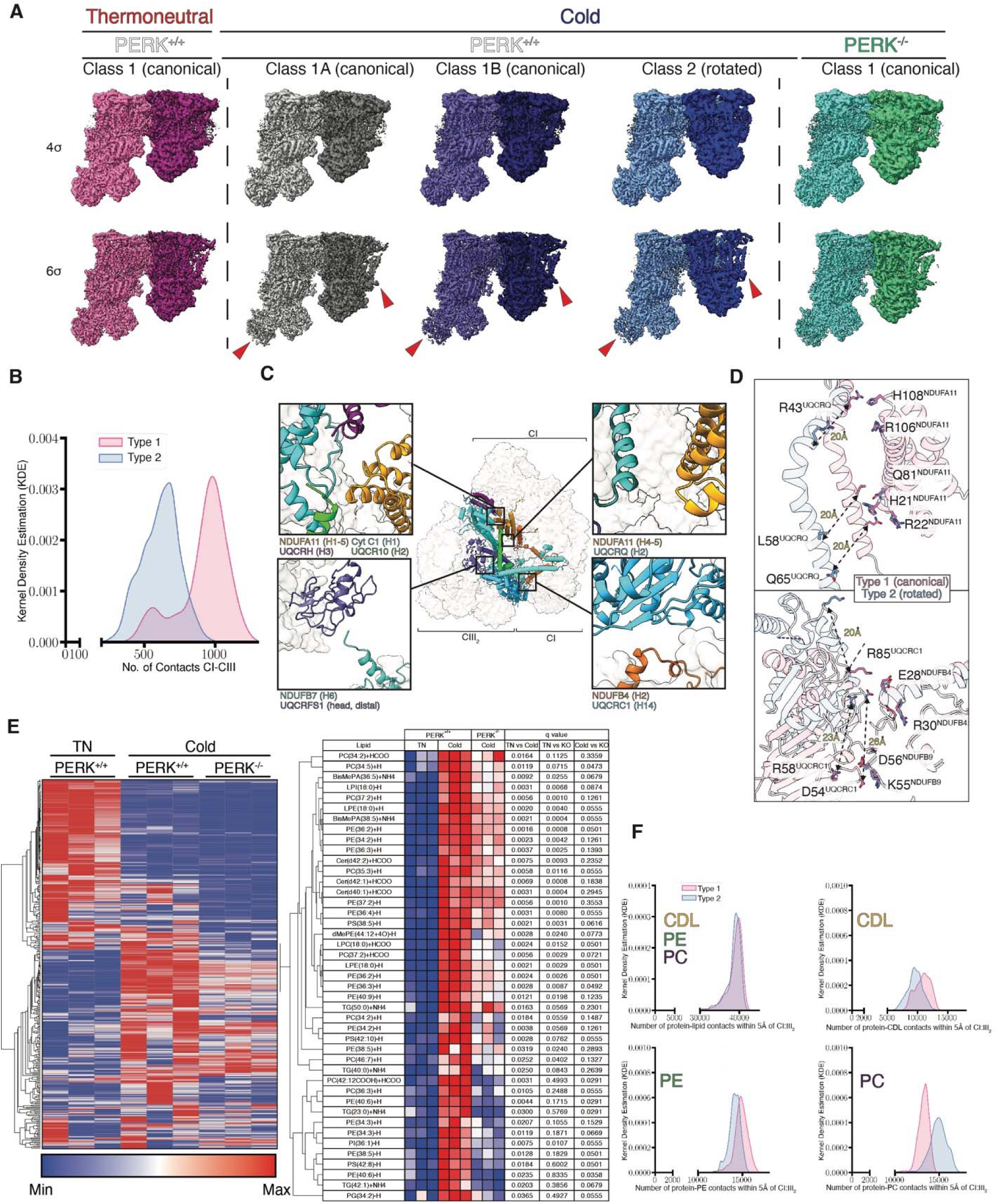
CI:III_2_ flexibility and activity. (**A**) EM density maps of identified classes represented at 4 and 6σ contour. Red arrows indicate flexible areas defined by EM maps. CIII_2_ densities are represented in darker colors. Type 1^WT-TN^, pink; type 1A^WT-cold^, grey; type 1B^WT-cold^, slate blue; type 2^WT-cold^, blue; type 1^KO-cold^, turquoise. (**B**) The kernel density estimation (KDE) of the number of protein-protein contacts of heavy atoms between CI and CIII_2_ with a 5 Å cutoff based on all simulations replicas of type 1 and type 2 assemblies. (C) Rotation of CIII_2_ complexes reorganizes the interaction landscape and chain-chain proximities in type 2 complexes. (**D**) Comparison of type 1 and 2 atomic models shows disruption of canonical CI:CIII_2_ interactions(17). Cα-Cα distances between pairs of interactions are shown. Side chains are used for depiction purposes. (**E**) Lipid composition in isolated mitochondria from WT (PERK^+/+^) mice under thermoneutral (TN) or cold adapted conditions and cold adapted PERK KO (PERK^-/-^) mice. (F) Quantitation of lipid contacts within 5 Å of the protein, from the simulations depicted in Figure 3F.

**Figure S8.**
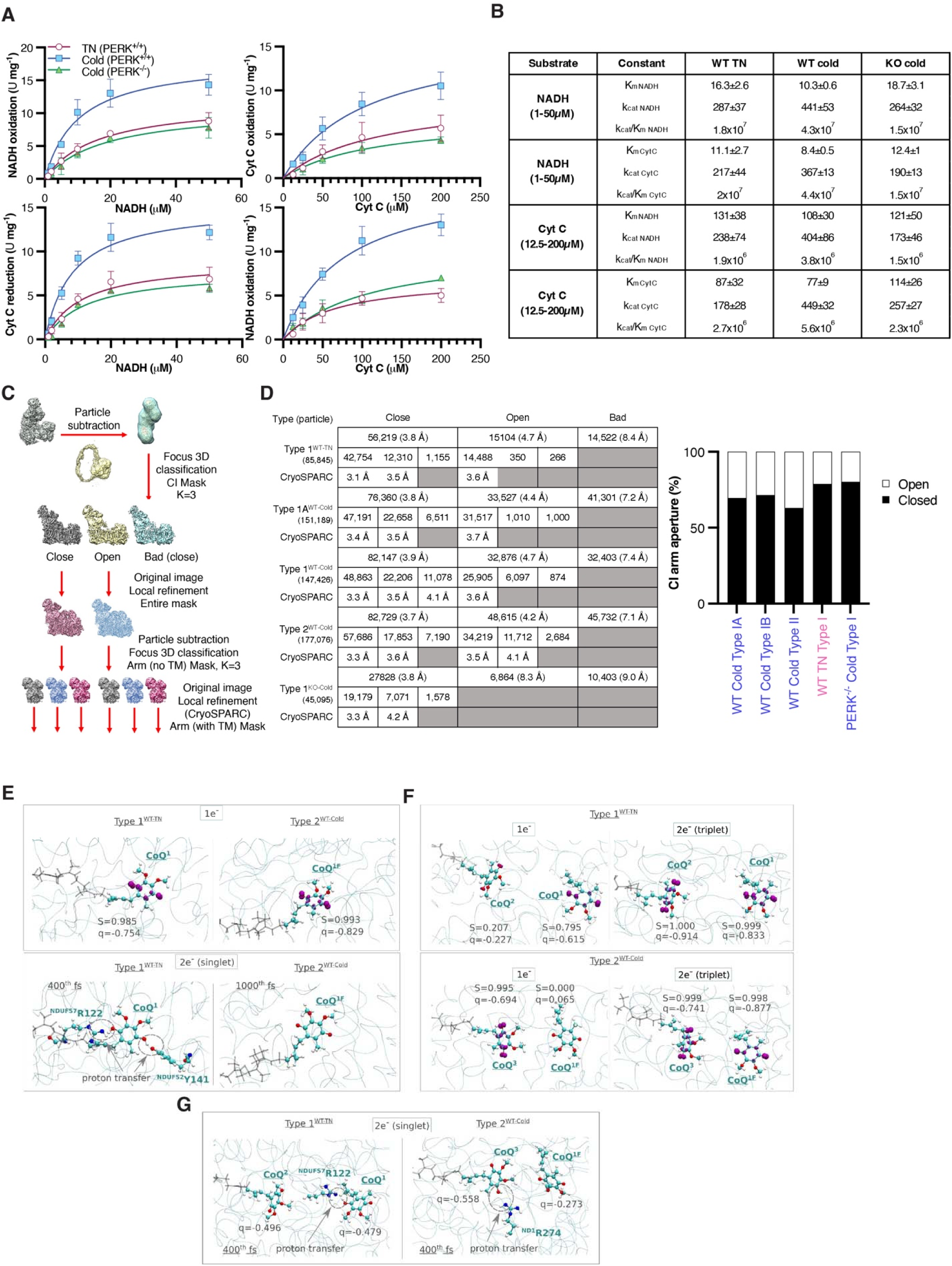
Determination of CI:III_2_ activities. (**A**) **Michaelis-Menten** kinetics of isolated CI:CIII_2_ complexes from iBAT exposed to different thermal conditions. TN; thermoneutral. N=3, n=3. (**B**) Apparent Michaelis-Menten parameters from the experiments represented in (A). (C and D) Focused 3D classification to define open and closed conformations of CI across different states. Graph on the right represents the proportion (%) of open and closed states across conditions. (E) QM/MM molecular dynamics simulation of single-quinone setups in type 1^WT-TN^ (left panel) and type 2^WT-Cold^ (right panel) complex I structures. Spin and charge population on CoQ_9_ after QM/MM molecular dynamics minimization upon one-electron reduction (above). The spin density is shown as a magenta isosurface (with an isovalue of 0.01), with captions indicating the total spin (S) and charge (q) population on CoQ species. Simulation snapshots from the unbiased QM/MM molecular dynamics trajectory upon two electron reduction (below). Superscripts (1 and 1^F^) on CoQ labels indicate the binding site locations. (F) QM/MM molecular dynamics simulations of the 2-quinone setups. Snapshots displayed are at the 400^th^ fs of the unbiased molecular dynamics trajectory from the type 1^WT-TN^ (top) and type 2^WT-Cold^ (bottom) structures. The one-electron reduced case is shown on the left and the two-electron case – on the right. The binding locations of CoQ ligands (sites 1, 1^F^, 2, and 3) are denoted in superscripts. Spin density populations are shown in magenta as isosurfaces with an isovalue of 0.01. Captions indicate the total spin (S) and charge (q) on the respective CoQ ligands. (G) QM/MM molecular dynamics simulations of the 2-quinone setups upon two electron reduction (singlet case) from type 1^WT-TN^ (left) and type 2^WT-Cold^ (right) structures. The snapshots are shown for the 400^th^ fs of the unbiased molecular dynamics trajectory. Superscripts denote the CoQ binding location in the quinone tunnel (sites 1, 1^F^, 2, and 3).

**Figure S9.**
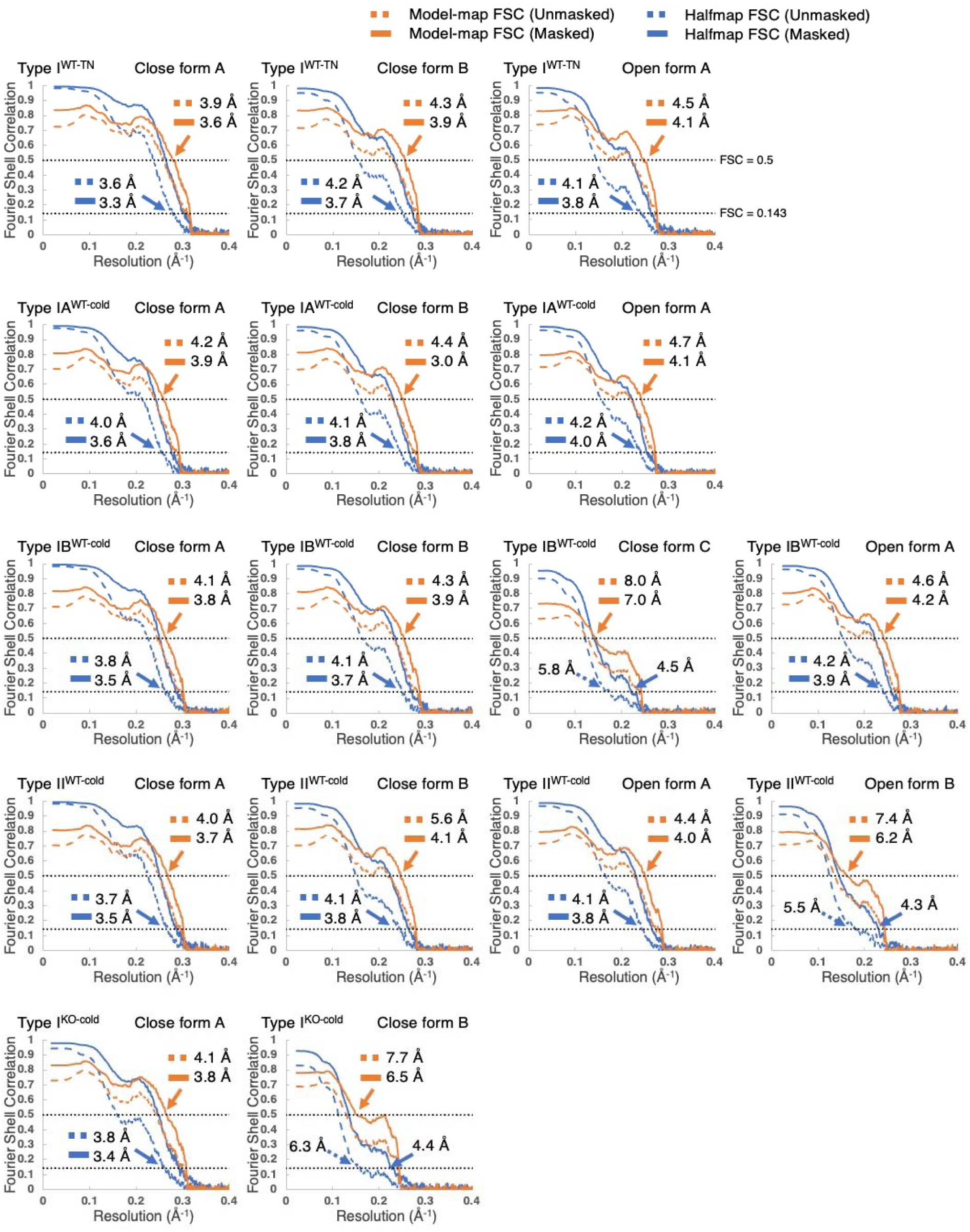
Fourier shell correlation (FSC) curves of CI peripheral arm across conditions. The gold standard half-map FSC and model-map FSC are shown as blue and orange lines, respectively (dashed line for unmasked and solid line for masked). The gold standard FSC cutoff (0.143) and model-map cutoff (0.5) are indicated with dashed black lines and estimated resolution on the cutoffs are indicated with colored arrows.

**Figure S10.**
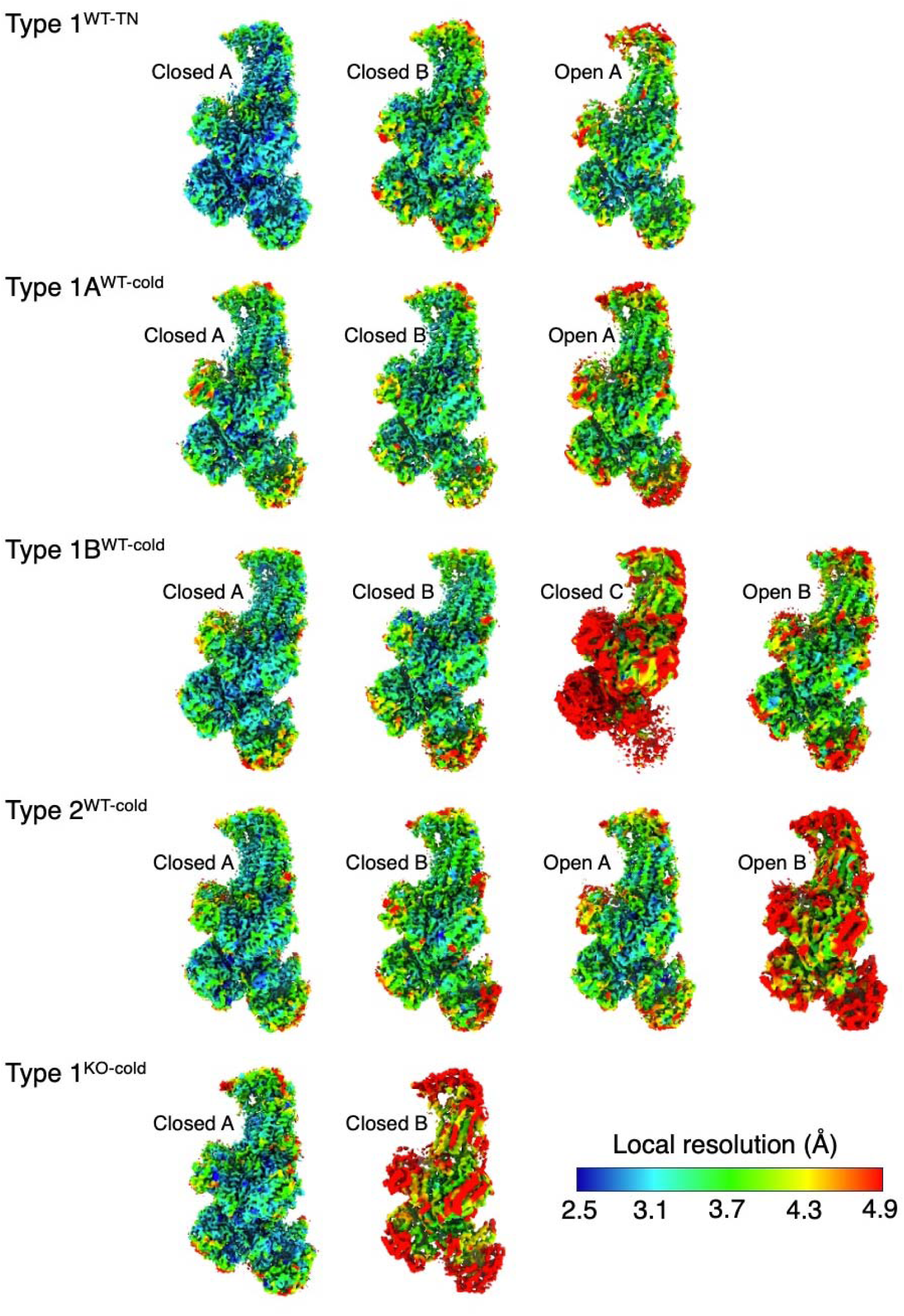
Local resolution estimates for CI peripheral arm across states. Determination of local resolution estimates for the focused refined CI peripheral arm maps in open and closed states with masks.

**Figure S11.**
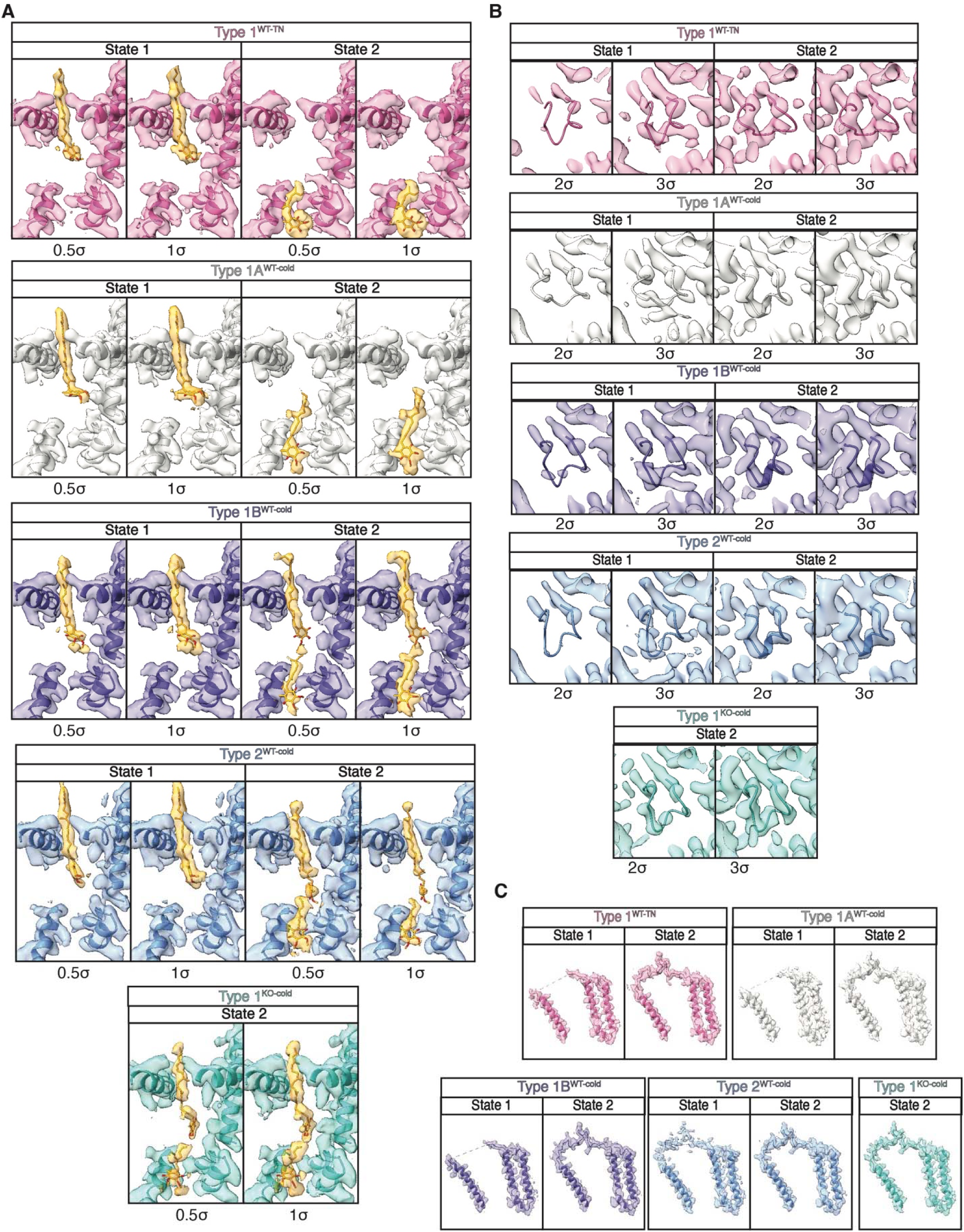
Presence of CoQ and ordering of NDUFS2 and ND3 loops in CI closed states. (**A**) EM map densities superimposed to atomic models to show CoQ presence at the Q chamber in CI. (**B**) Superimposition of Cryo-EM densities to NDUFS2 loop from atomic models at 2 and 3σ contour. (**C**) Superimposition of Cryo-EM densities to ND3 loop from atomic models at 1σ contour. In (**A-C**), Type 1^WT-TN^, pink; type 1A^WT-cold^, grey; type 1B^WT-cold^, slate blue; type 2^WT-cold^, blue; type 1^KO-cold^, turquoise. Red triangles point at the indicated loops on the right.

**Figure S12.**
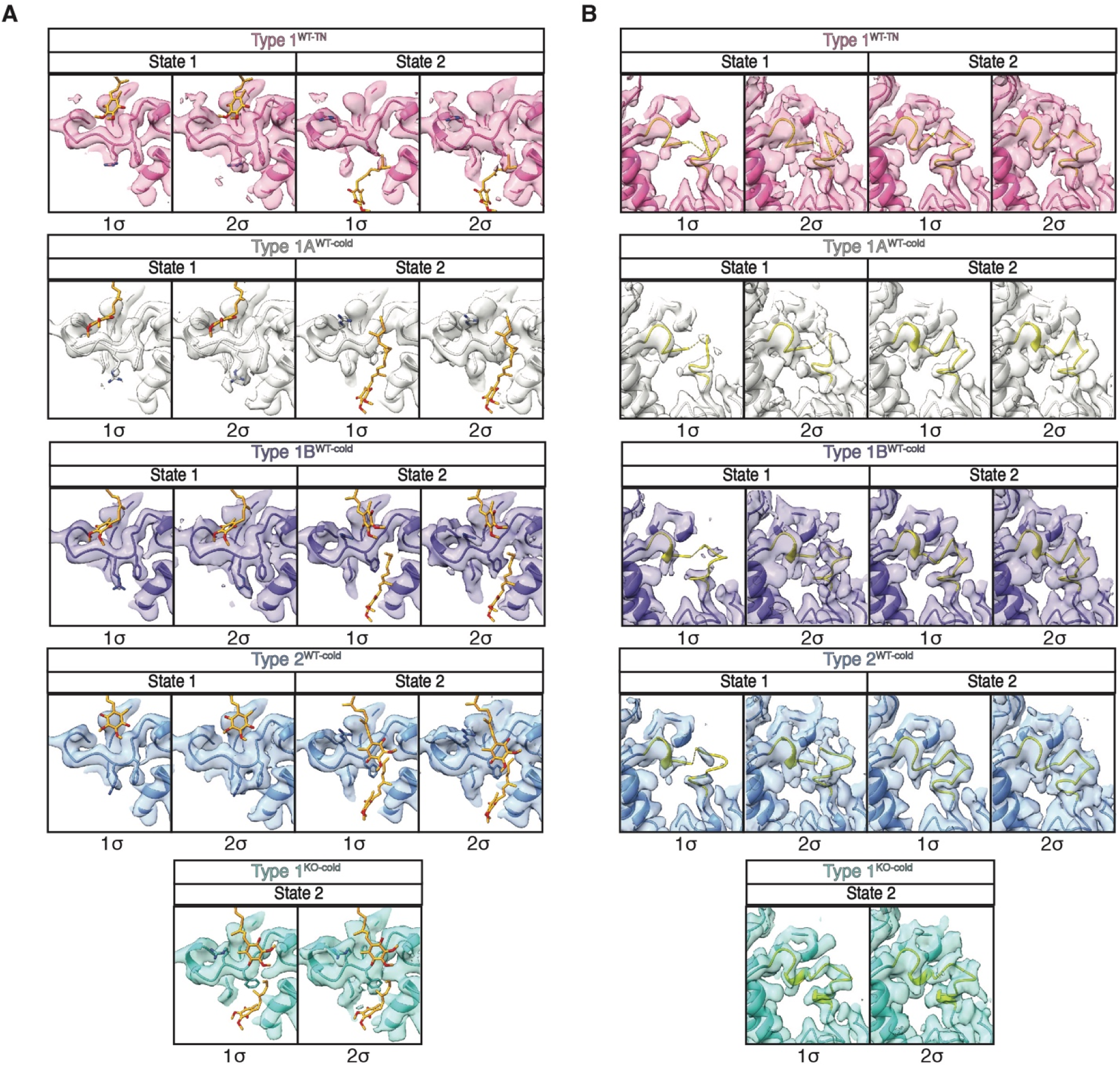
Ordering of NDUFS8 and ND1 loops in CI closed states. (**A**) Superimposition of Cryo-EM densities to NDUFS8 loop from atomic models at 1 and 2σ contour. Flipping of residues F121 and R122 is represented along with CoQ_9_ (orange). (**B**) Superimposition of Cryo-EM densities to ND1 loop (yellow in atomic model) from atomic models at 1 and 2σ contour. In (**A** and **B**), Type 1^WT-TN^, pink; type 1A^WT-cold^, grey; type 1B^WT-cold^, slate blue; type 2^WT-cold^, blue; type 1^KO-cold^, turquoise. Red triangles point at the indicated loops on the right.

**Figure S13.**
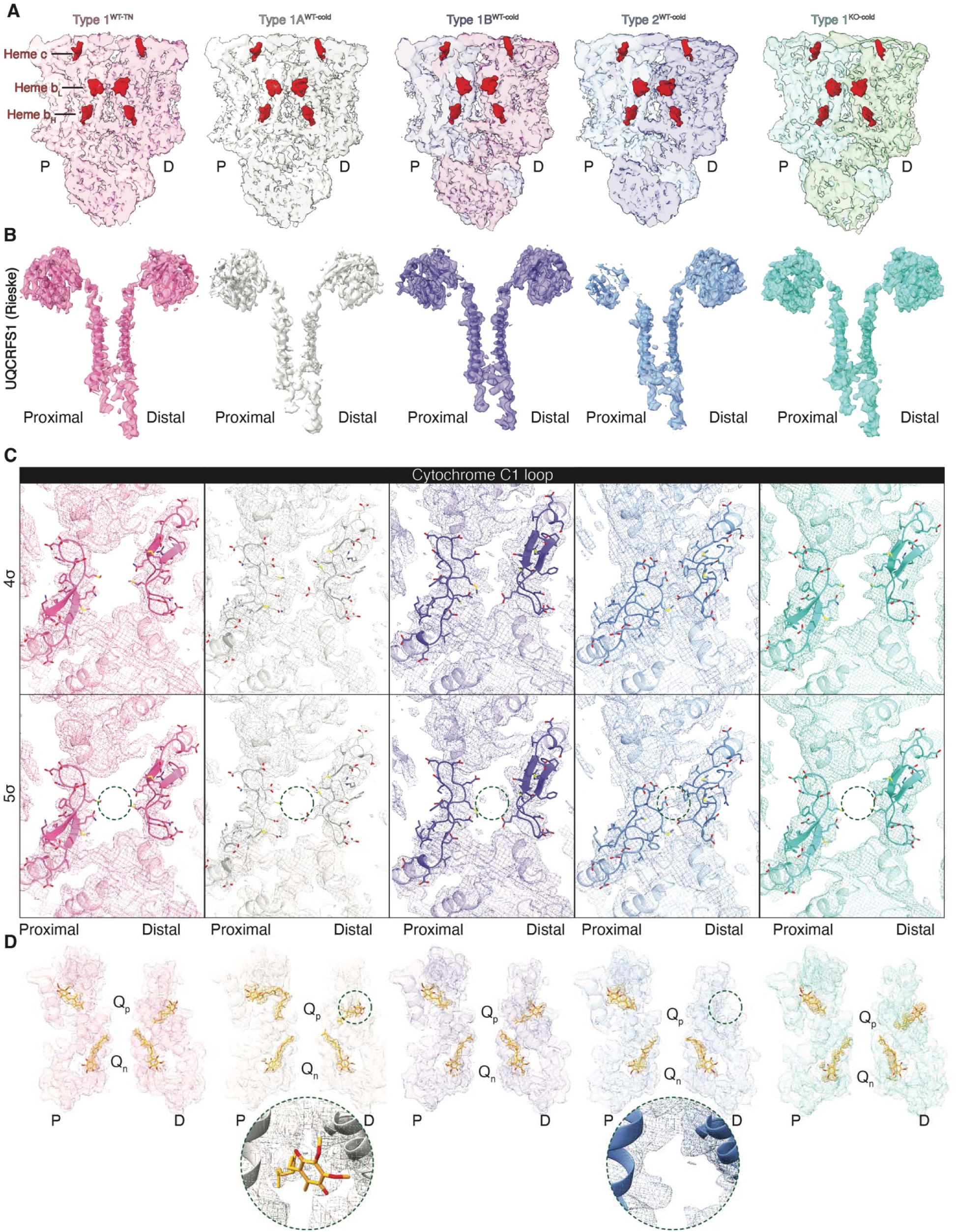
Comparison of CIII_2_ structure across states. (**A**) Cross-sectional representation of CIII_2_ complexes across states shows symmetry preservation. P; proximal to CI. D; distal to CI. (**B**) Superimposition of UQCRFS1 (Rieske) EM densities to atomic model (3σ contour) shows substantial density decrease at the head and neck of the proximal subunit. (**C**) Superimposition of EM maps (4 and 5σ contour) to atomic models shows high flexibility of the Cyt. C loop (residues 150-170) and approximation to its counterpart in the other CIII protomer in type 2 complexes. Side chains are shown to depict the acidic composition of the loop. (**D**) Superimposition of EM maps (3σ contour) to atomic models shows potential distribution of CoQ molecules in CIII_2_. Type 2 complexes show no occupancy of the distal oxidative site. P; proximal. D; distal. Q_p_; oxidative site. Qn; reductive site. In (**A-D**), Proximal = close to CI active site. Distal = further from CI active site. Type 1^WT-TN^, pink; type 1A^WT-cold^, grey; type 1B^WT-cold^, slate blue; type 2^WT-cold^, blue; type 1^KO-cold^, turquoise.

**Table S1.** Cryo-EM data collection.

| **Data collection** | **Wt-thermoneutral** | **Wt-cold acclimated** | **PERK KO-cold acclimated** |
| --- | --- | --- | --- |
| Microscope | UMass Talos Arctica | UMass Talos Arctica | Harvard Titan Krios |
| Detector | K3 | K3 | K3 |
| Magnification | 36,000 | 36,000 | 105,000 |
| Voltage (kV) | 200 | 200 | 300 |
| Electron Exposure (e<sup>-</sup>/Å<sup>2</sup>) | 46.6 | 46.1/45.9 | 50.2 |
| Dose per frame (e<sup>-</sup>/Å<sup>2</sup>) | 1.261 | 1.536/1.509 | 1.226 |
| Defocus Range (µm) | -1.0 to -2.2 | -1.0 to -2.2 | -1.0 to -2.0 |
| Pixel Size (Å) | 1.1 | 1.1 | 0.83 |
| Acquisition software | SerialEM | SerialEM | SerialEM |
| Collected micrograph (No.) | 5,262 | 3,294/7,004 | 5,220 |

**Table S2.**
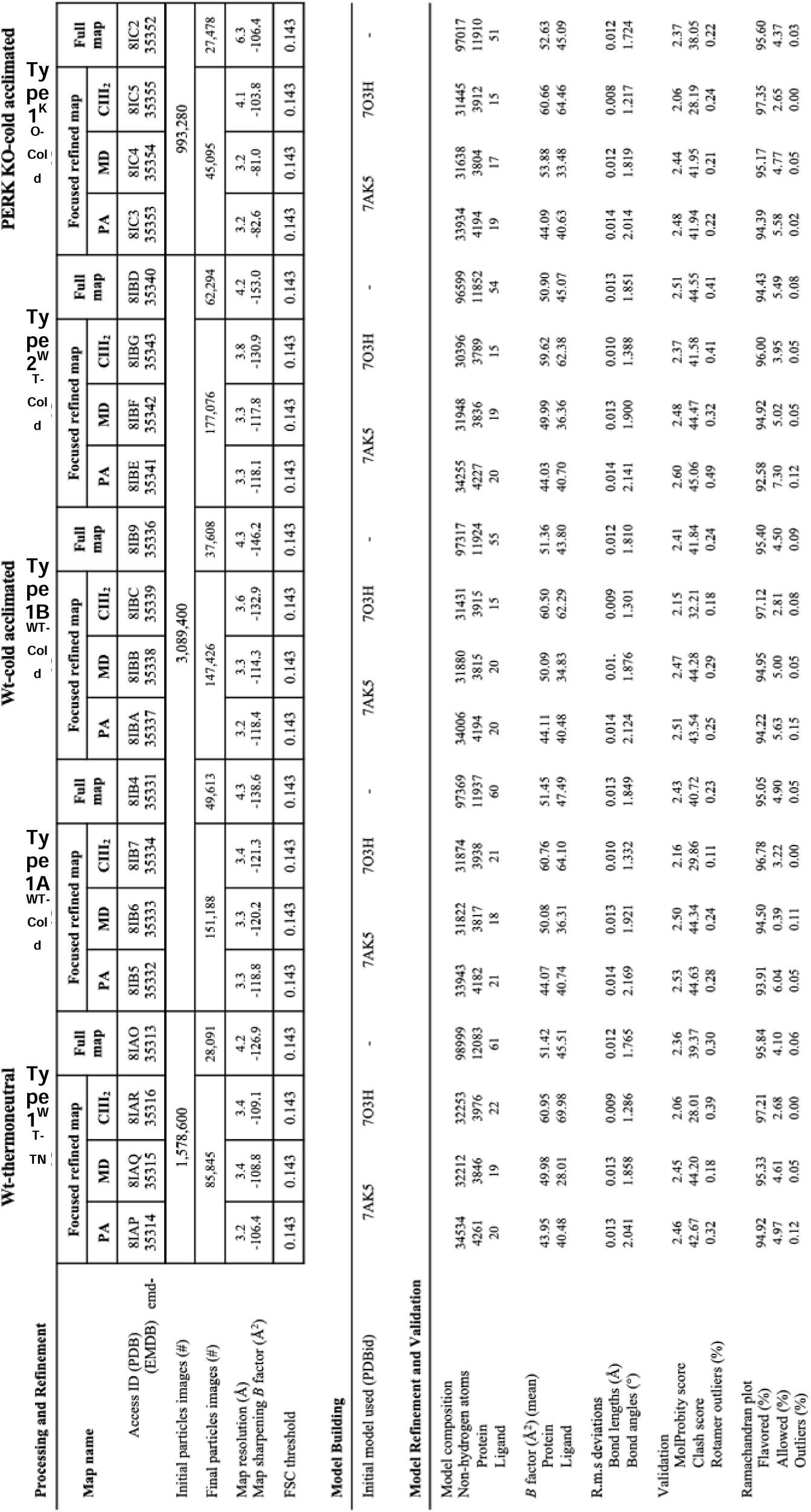
Cryo-EM data processing, map and model validation.

**Table S4.**
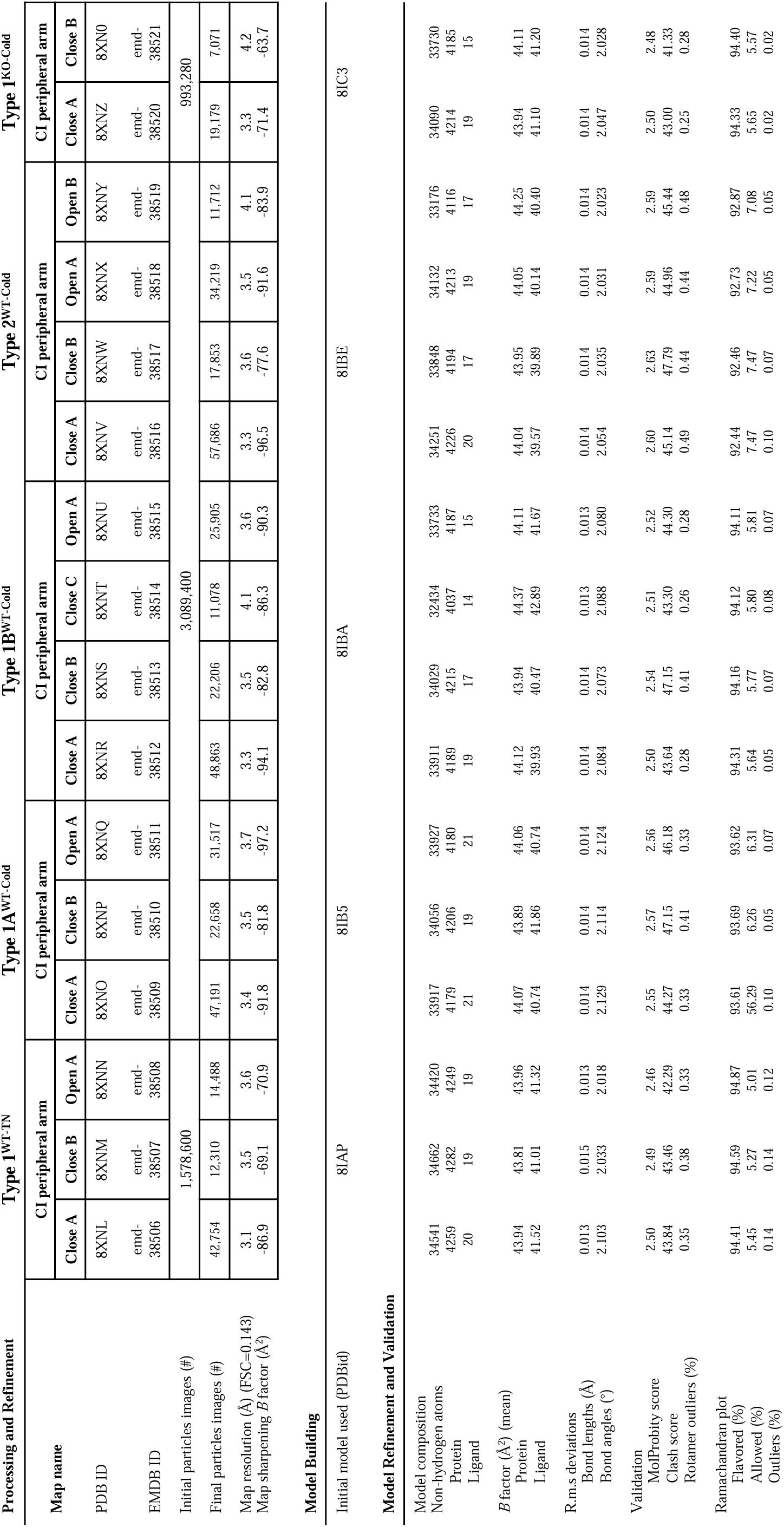
Cryo-EM data processing, map and model validation for CI peripheral arm.

**Table S5.** Positions of CoQ species modelled in different sites of the quinone binding cavity in respiratory complex I.

| Setup | Quinone type | Structure |  |
| --- | --- | --- | --- |
|  |  | Type 1<sup>WT-TN</sup> | Type 2<sup>WT-cold</sup> |
| 1-quinone setups (1Q) | CoQ<sub>9</sub> | Site 1 | Site 1F |
| 2-quinone setups (2Q) | CoQ<sub>1</sub> | Site 1 | Site 1F |
|  | CoQ<sub>9</sub> | Site 2 | Site 3 |

**Table S6.** QM regions considered in our calculations.

| Setup | Structure | Number of atoms<br>(excluding link<br>hydrogens) | Residues included in the QM region |
| --- | --- | --- | --- |
| 1-quinone<br>setups | Type 1<sup>WT-TN</sup> | 133 | NDUFS7<sup>Ala101</sup>, NDUFS7<sup>Val102</sup>, NDUFS7<sup>Met105</sup>,<br>NDUFS7<sup>Arg122</sup>, NDUFS2<sup>Hsd92</sup>, NDUFS2<sup>Tyr141</sup>,<br>NDUFS2<sup>Met185</sup>, NDUFS2<sup>Thr189</sup>, NDUFS2<sup>Asp193</sup>,<br>NDUFS2<sup>Val457</sup>, **CoQ<sub>9</sub>** |
|  | Type 2<sup>WT-Cold</sup> | 144 | NDUFS7<sup>Thr94</sup>, NDUFS7<sup>Ala101</sup>, NDUFS7<sup>Met104</sup>,<br>NDUFS7<sup>Met105</sup>, NDUFS7<sup>Phe121</sup>, NDUFS7<sup>Arg122</sup>,<br>NDUFS2<sup>Thr189</sup>, NDUFS2<sup>Leu192</sup>, NDUFS2<sup>Asp193</sup>,<br>ND1<sup>Glu204</sup>, ND1<sup>Ser205</sup>, **CoQ<sub>1</sub>**, **CoQ<sub>9</sub>** |
| 2-quinone<br>setups | Type 1<sup>WT-TN</sup> | 244 | NDUFS7<sup>Trp91</sup>, NDUFS7<sup>Met105</sup>, NDUFS7<sup>Arg122</sup>,<br>NDUFS7<sup>Ser124</sup>, NDUFS7<sup>Gln127</sup>, NDUFS2<sup>His92</sup>,<br>NDUFS2<sup>Tyr141</sup>, NDUFS2<sup>Met185</sup>, NDUFS2<sup>Thr189</sup>,<br>NDUFS2<sup>Asp193</sup>, ND1<sup>Asp199</sup>, ND1<sup>Glu202</sup>,<br>ND1<sup>Glu204</sup>, ND1<sup>Asn212</sup>, ND1<sup>Phe220</sup>, **CoQ<sub>9</sub>** |
|  | Type 2<sup>WT-Cold</sup> | 257 | NDUFS7<sup>Trp91</sup>, NDUFS7<sup>Met105</sup>, NDUFS7<sup>Phe121</sup>,<br>NDUFS7<sup>Ser124</sup>, NDUFS7<sup>Gln127</sup>, NDUFS2<sup>Thr189</sup>,<br>NDUFS2<sup>Leu192</sup>, NDUFS2<sup>Asp193</sup>, ND1<sup>Glu24</sup>,<br>ND1<sup>Arg25</sup>, ND1<sup>Glu202</sup>, ND1<sup>Glu204</sup>, ND1<sup>Ser205</sup>,<br>ND1<sup>Tyr228</sup>, ND1<sup>Phe220</sup>, ND1<sup>Arg274</sup>, **CoQ<sub>1</sub>**, **CoQ<sub>9</sub>** |

**Table S7.** Electron transfer rate constants (k_ET_), estimated according to the Moser-Dutton ruler (formula (1), “Computational methods”). Edge-to-edge distance between the donor and acceptor is calculated from the structure.

| Setup | Donor | Acceptor | Structure | R, Å | Edge-to-edge distance between the atoms | $k_{ET}, *10^3 \text{ 1/s}$ |
| --- | --- | --- | --- | --- | --- | --- |
| 1Q | N2 | CoQ<sub>9</sub> | Type 1<sup>WT-TN</sup> | 11.20 | S2(F4R) - C1(Q9) | 5370.32 |
|  |  |  | Type 2<sup>WT-cold</sup> | 14.62 | S4(F4R) - O4(Q9) | 47.64 |
| 2Q | CoQ<sub>1</sub> | CoQ<sub>9</sub> | Type 1<sup>WT-TN</sup> | 15.81 | O2(Q9) - O5 (Q1) | 9.20 |
|  |  |  | Type 2<sup>WT-cold</sup> | 8.60 | O5(Q1) - O4(Q9) | 194984.46 |

**Movie S1.** Comparison between type 1 and type 2 CI:III_2_ complexes.

## Acknowledgements

NIH RO1 DK081418 and RO1 DK089883 (PP). Human Frontier Science Program LT000033/2019-L (PLM). NIH NIDDK K99DK133502 (PLM). Deutsche Forschungsgemeinschaft (DFG): Projektnummer 501493132 (NB). Academy of Finland, the Sigrid Juselius Foundation, the Jane and Aatos Erkko Foundation and the Magnus Ehrnrooth Foundation (VS). CSC (Center for Scientific Computing, Finland) (VS). SUSTech Institute for Biological Electron Microscopy (ML).

## Author contributions

Conceptualization: PP, ML and PLM. Methodology: YCS, PLM, AD, OZ and VS. Investigation: YCS, PLM, AD, OZ, CFB, NB, KS and CX. Visualization: PLM, OZ and AD. Funding acquisition: PP, PLM and VS. Supervision: PP, ML, PLM and VS. Writing – original draft: PLM. Writing – review & editing: YCS, PLM, AD, OZ, CFB, NB, VS, ML and PP.

## Data and materials availability

All data will be available upon reasonable request to the corresponding authors. Maps and atomic models are available at the EMDB and RCSB. PDB ID: type I^WT-TN^ (8IAP, 8IAQ, 8IAR, 8IAO, 8XNL, 8XNM, 8XNN), type IA^WT-cold^ (8IB5, 8IB6, 8IB7, 8IB4, 8XNO, 8XNP, 8XNQ), type IB^WT-cold^ (8IBA, 8IBB, 8IBC, 8IB9, 8XNR, 8XNS, 8XNT, 8XNU), type II^WT-cold^ (8IBE, 8IBF, 8IBG, 8IBD, 8XNV, 8XNW, 8XNX, 8XNY), type I^KO-cold^ (8IC3, 8IC4, 8IC5, 8IC2, 8XNZ, 8XN0), EMDB ID: type I^WT-TN^ (emd-35314, emd-35315, emd-35316, emd-35313, emd-38506, emd-38507, emd-38508), type IA^WT-cold^ (emd-35332, emd-35333, emd-35334, emd-35331, emd-38509, emd-38510, emd-38511), type IB^WT-cold^ (emd-35337, emd-35338, emd-35339, emd-35336, emd-38512, emd-38513, emd-38514, emd-38515), type II^WT-cold^ (emd-35341, emd-35342, emd-35343, emd-35340, emd-38516, emd-38517, emd-38518, emd-38519), type I^KO-cold^ (emd-35353, emd-35354, emd-35355, emd-35352, emd-38520, emd-38521).

## Materials and Methods

### Animal breeding

Animals were generated as previously described(2). Briefly, PERK^flox/flox^ mice were from Jackson Labs (stock 023066)(47) and crossed with adiponectin Cre-expressing (stock 010803)(48) mice to generate adipocyte specific PERK^-/-^ mice. Cre negative littermate controls (PERK^+/+^) were used as wild-type mice(48). Animals were housed under room temperature 22°C conditions and 12/12 h light/dark cycles with free access to food and water. 12 weeks old males were transferred to either cold (4°C) or thermoneutral (28°C) conditions for 8 days prior to tissue isolation. All the procedures in this paper were performed in compliance with the 1ACUC approved protocols at the Animal Research Facility (Beth Israel Deaconess Medical Center, Boston, MA, USA).

### Complex isolation

Isolation was performed as previously described with minor modifications(2). Interscapular brown adipose (iBAT) were obtained from 16 mice each condition and dounce-homogenized in 25 mM HEPES, pH 7.5, 225 mannitol, 75 mM sucrose, 0.1 mM EGTA and protease and phosphatase inhibitors until the solution was homogeneous in color and texture (10 strokes). Samples were centrifuged at 800 g and supernatant was collected. Pellet containing nuclei was resuspended and homogenized in isolation buffer and centrifuged again to collect mitochondria in the supernatant. Mitochondria were pelleted at 7500 g for 10 min in the cold and separated by ultracentrifugation using a Percoll gradient(49). Mitochondria were collected at the bottom of the ultracentrifuge tube and diluted ten times in 5 mM HEPES, pH 7.4, 250 mM mannitol, 0.1 mM EDTA and pelleted at 9000 g for 10 min in the cold. Mitochondria were resuspended in isotonic sucrose buffer (30 mM HEPES, pH 8.0, 250 mM sucrose, 0.1 mM EDTA) and sonicated four times, 5 seconds each, 100 watts (20% amplitude) and 30 seconds off between sonication cycles. Lysates were centrifuged at 9000 g for 10 minutes and supernatant was collected and centrifuged at 200,000 g in isotonic sucrose buffer. Membranes were collected at the bottom and solubilized in the cold in 30 mM HEPES, pH 8.0, 10% glycerol, 150 mM NaCl, 0.1 mM EDTA, 1% digitonin and protease inhibitors for 2 hours. Samples were centrifuged at 18000 g and supernatant injected in a Superose 6 10/300 GL column (Cytiva) attached to an ÄKTA Pure 25 (Cytiva) equilibrated in 30 mM HEPES, pH 8.0, 150 mM NaCl, 0.1 mM EDTA, 0.02% GDN. Fractions were collected, flash-frozen in liquid nitrogen and kept at –80°C.

### Determination of complex activity

Complex I:III_2_ activity assays were performed as described(50) using flat bottom low absorbance 96 well-plates (Corning). Mixtures contained 25 mM K-phosphate buffer, pH 7.4, 1 mg/mL fatty acid free BSA (GoldBio), 50 μM oxidized Cyt C (Sigma), 30 μM KCN (Sigma), and 5-10 nM of protein complexes in a final volume of 40 μL. Reactions started by addition of 0.2 mM NADH (Sigma). Plates were shaken for 20 seconds before the assay, and absorbances at 340 (NADH oxidation) and 550 nm (Cyt C reduction) were recorded simultaneously at 30°C using a Clariostar plus plate reader (BMG Lab Tech). Activities in the presence of CI inhibitor rotenone (10 μM) (Sigma) were used as blank and calculated using ε_NADH_=6.22 mM^-1^cm^-1^ and ε_Cyt C_=18.5 mM^-1^ cm^-1^.

### Negative-stain EM Screening for respiratory complex from iBAT mitochondria

Different fractions from size exclusive chromatography (SEC) were examined using negative-stain EM for identification of respiratory complex. All fractions were individually prepared to be in the concentration range of 0.02 – 0.06 mg/ml, applied on glow discharged EM grids (G400, Ted Pella, Inc.) covered with a thin layer of continuous carbon film, and stained with 1.5% (w/v) uranyl formate. EM grids were imaged using a Philips CM10 electron microscope (Thermo Fisher Scientific), operated at 100 kV and at nominal magnification of 52,000x. Negative-stain EM images were analyzed using SamViewer(51), an interactive image analysis program.

### Cryo-EM sample preparation and data collection

The SEC fractions containing respiratory complexes were combined and concentrated to 0.33 mg/ml. 2.5 – 3.5 μL sample was applied to graphene oxide coated EM grids (Quantifoil Cu400 R1.2/1.3) following a previously published protocol(52). Sample was kept on the grid for 5 s, blotted for 2.5 – 3 s at room temperature with 85% humidity using a Gatan Cryoplunge 3 system, and vitrified in liquid ethane cooled by liquid nitrogen. Cryo-EM images of cold acclimated and thermoneutral samples were collected on a Talos Arctica (Thermo Fisher Scientific), and those from PERK^-/-^ mouse samples were collected on a Titan Krios (Thermo Fisher Scientific). The detailed information and parameters of data collection are listed in Table S1.

### Cryo-EM data processing

Detailed information of number of particles and map resolutions are described in Table S2. Movies were motion corrected with MotionCor2(53), and defocus values calculated with CTFFIND4(54). All other 2D image processing were performed using Simplified Application Managing Utilities of EM Labs (SAMUEL)(51). Briefly, dose-weighted images were binned 6x (for cold acclimated and thermoneutral samples) or 8x (for PERK^-/-^ sample) with “sampilcopy2d.py” and interactively screened. Particle picking, particle stack generation, and 2D classification were performed with “samautopick.py”, “sampilboxparticle.py” and “samtree2dv3.py”, respectively. Additional 2D and 3D classification steps were performed to exclude bad particles using RELION 2.0(55) and RELION 3.0(56), respectively. After sorting the 6x binned particles stack, 3x binned particle stacks (bin4 for cold PERK^-/-^) were generated for further 3D classifications. First round of 3D classification was carried out with K=6 for 35 cycles in RELION 3.0 and good classes (one class from TN WT, three classes from cold WT and one class from cold PERK^-/-^) were selected for second round 3D classification after keeping all particles assigned in those selected classes within the final 5 cycles. After selecting particles from second round 3D classification, we performed auto-refinement with original, unbinned particle stack using the parameters of 30-Å lowpass filter for reference, 7.5 degree for angular sampling and 1.8 degree for starting local searches. Then, map sharpening was performed by using “relion_postprocess” and yielded 3.9 – 4.1 Å in all conformations but the densities corresponding to CIII_2_ position were less well defined.

While most classes of supercomplex were determined to high resolution after auto-refinement, CIII_2_ complex showed lower resolution relative to the transmembrane domains of CI complex, likely due to minor movement of CIII_2_ along the transmembrane domains of CI. For determining the stable position of CIII_2_, masked 3D classification focusing on CIII_2_ with signal subtraction of CI was performed for all conformations. After running 3D classification without alignment (K=3, 50 cycles, T value =20 and CIII_2_ mask), we selected the best class with ordered transmembrane domain density, followed by local refinement using original images and the full mask for the entire consensus map. For obtaining stable ‘full map’ of supercomplex, parameters of angular sampling and local searches from a sampling were changed to 3.7 degree. Resolution estimation using “relion_postprocess” yielded 4.2 – 4.3 Å (type 1^WT-TN^, types 1A^WT-cold^, 1B^WT-cold^ and 2^WT-cold^) and 6.3 Å (I^KO-cold^) from overall mask and 4.4 Å (type 1^WT-TN^ and types 1A^WT-cold^), 6.1 Å (type 1B^WT-cold^), 6.4 Å (type 2^WT-cold^), and 8.6 Å (I^KO-cold^) from CIII_2_ mask, respectively. The resolutions of full maps were estimated based on the FSC=0.143 criterion and curves were calculated using “relion_postprocess” (Figure S2).

To push higher resolution of each region, further improved maps were obtained by focused refinement using several different maskS6^3^. In order to generate reference maps and masks, the structural models of mouse CI (7AK5)(23) and CIII_2_ (7O3H)(14) were individually fitted in the stable ‘full map’ using “fit in map” in UCSF Chimera, and the models corresponding to peripheral arm, membrane domain, and CIII_2_ were extracted by PyMOL (Version 2.5.2, Schrödinger, LLC.). These extracted models were converted to 20 Å-resolution maps using “molmap” in UCSF Chimera, and masks were generated from these maps using “relion_mask create” in RELION with 2 pixels extension and 6 pixels for smoothening of the edges. “Local refinement” in cryoSPARC(57) with these maps and masks was performed for calculating the focused refined maps, resulting in 3.2 Å – 3.4 Å resolutions for peripheral arm and membrane domain, and 3.4 Å – 4.1 Å resolution for CIII_2_ (type 1^WT-TN^ 3.4 Å, types 1A^WT-cold^ 3.4 Å, type 1B^WT-cold^ 3.6 Å, type 2^WT-cold^ 3.8 Å and type 1^KO-cold^ 4.1 Å, respectively). Fourier shell correlation (FSC) curves of focused refined maps were computed using “Comprehensive Validation” in Phenix (Figure S3).

### Model building, refinement, and display

We built three models which fit in the map of peripheral arm, membrane domain and CIII_2_. Mouse CI (7AK5)(23) and CIII_2_ (7O3H)(14) were used as the starting templates for model building. Each subunit in templates was individually fitted to the focused refined maps using “fit in map” function in UCSF chimera and the model refined using “Real space refine zone” in Coot(58). Side chain rotamers, varied secondary structures, and bound ligands were further refined manually at the reasonable sigma levels (3.0∼7.0). The transmembrane helices around the distal lobe of CIII_2_, those with weak densities in type 2^WT-cold^ (CJ/J/ and CK/K/) and type 1^KO-cold^ (CK/K/), were replaced with well-built models from types 1A^WT-cold^ and type 1^WT-TN^, respectively. Final models built from focused refined maps were validated using “Comprehensive validation” in Phenix(59), and fitted in the stable ‘full map’ of entire supercomplex. Final entire models of respiratory supercomplex were used as a reference model for the alignment of focused refined maps, and these maps were combined together using “combined focused maps” function in Phenix. Figures of structural models, cryo-EM maps and combined focused maps were generated using UCSF Chimera, ChimeraX(60). and PyMOL. Model quality statistics were calculated from Comprehensive Validation (Cryo-EM) in Phenix. <u>Lipidomics</u>

Mitochondria were isolated as previously described (see above)(49). Lipids were extracted as previously indicated(25). Briefly, 100 μg of pure mitochondria were resuspended in 200 μl of H_2_O (MS-grade) and mixed with 1.5 ml of HPLC-grade methanol and 5 ml of methyl tert-butyl ether (MTBE) and mixed for 1h. Phase separation was achieved by adding 1.25 ml of H_2_O and centrifugation at 1,000g for 10 min. The upper MTBE phase was collected vacuum dried at 4 °C. Samples were resuspended in 30 μl of 1:1 LC–MS grade isopropanol:acetonitrile methanol and subjected to lipid LC-MS/MS analysis using the QExactive Plus Orbitrap (ThermoFisher Scientific) instrument. Data analysis was performed as follows. Identical lipid entries were consolidated and lipids with missing values discarded. Two-way ANOVA analyses and multiple testing FDR comparisons were used with a 0.05 significance threshold.

### Atomistic MD simulations of CICIII_2_ supercomplexes

Classical atomistic molecular dynamics simulations were conducted using the type 1^WT-TN^ and type 2^WT-Cold^ structures of supercomplexes. To address the issue of missing residues due to unresolved regions, we applied Modeller software(61). We used PDBs 7B93(62) for CI and 7O3H(14) for CIII as reference structures. It has recently been suggested that atomistic simulations of membrane protein structures when performed in protonation states pre-determined with pKa calculations are overall stable compared to when simulations performed with all amino acids in standard states(63). Therefore, protonation states of all titratable residues at pH 7 were assigned based on the pKa calculations (see below).

The model systems incorporated all lipids and other molecules resolved in the structures. This composite system was then embedded in a membrane that had been previously generated with the CHARMM-GUI platform(*64*). The membrane composition is 50% palmitoyl-oleoyl-phosphatidylcholine (POPC), 35% palmitoyl-oleoyl-phosphatidylethanolamine (POPE), and 15% cardiolipin, approximating the inner mitochondrial membrane composition. The structures were aligned to the lipid membrane through the utilization of OPM alignment(*65*). The solvent was composed of TIP3P(*66*) water molecules, along with 0.1 M of Na^+^ and Cl^-^ ions.

The force field parameters employed for various components of the system were uniformly based on the CHARMM36 force field(*67*, *68*), as well as oxidized quinones(*68*), FMN(*69*), FeS clusters(*70*), hemes(*71*), and NADPH(*72*).

After system construction, modeled water, lipids and ions that were overlapping with the protein were removed. Before the production run, the system underwent a series of two energy minimization and two equilibration stages. The initial energy minimization step, consisting of 10,000 steps, was executed using the NAMD software(*73*), imposing constraints solely on the heavy atoms of resolved structure, except for the missing regions that were modeled with homology modeling. Subsequently, in the second energy minimization phase, the constraints remained largely similar, with the additional constraints imposed on the phosphorous atoms of the membrane molecules in their z-coordinates. This stage was executed with GROMACS(*74*) and continued until the system achieved a force average of 1000 kJmol^-1^nm^-1^.

Following energy minimization, the system underwent an equilibration period of 10 nanoseconds in the NPT ensemble, maintaining the earlier imposed constraints. This process was repeated for an additional 10 nanoseconds, albeit with constraints solely applied to the backbone of the resolved protein segments. The latter step allowed settling of lipid membrane around the protein. Both equilibration stages employed the V-rescale thermostat(*75*) and the Berendsen barostat(*76*).

For the production runs, the system was subjected to a Nose-Hoover thermostat(*77*, *78*) and the Parrinello-Rahman barostat(*79*, *80*) for temperature and pressure control, respectively. All simulations were executed at a temperature of 310 K and a pressure of 1 atm employing a time step of 2 femtoseconds using LINCS(*80*) algorithm and long-range electrostatics dealt with Particle-mesh Ewald methods(*81*, *82*).

Each ∼1 s simulation (of type 1 and type 2) was repeated three times, resulting in a total of 6 s of simulation data. Quantitative and visual analyses were performed using VMD software(*82*).

### QM/MM simulations

Hybrid quantum-mechanical/molecular mechanical (QM/MM) molecular dynamics simulations were performed on the two cryo-EM structures of respiratory complex I from type 1^WT-TN^ and type 2^WT-cold^ conditions. Each model system was restricted to 6 core subunits of complex I (ND1, ND3, NDUFS2, NDUFS3, NDUFS8, and NDUFS9). The N terminus of NDUFS2 (residues 33 to 77) was excluded to prevent large fluctuations of this region in the pruned system. It has remained unclear if the observed densities in the Q-tunnel of complex I are of two different Q molecules occupying the tunnel at the same time or the same Q molecule at two different time points (Figure S9 and S11). Therefore, we created two QM/MM setups, one with a single CoQ_9_, and another with two CoQ molecules modeled at the Q_d_ and Q_s_ binding site regions(15). Since, the location of the tail of CoQ_9_ bound at Q_d_ sites is unclear, it was modeled as CoQ_1_. In the single-quinone (1Q) setups, the head group of CoQ_9_ was modelled at sites 1 and 1F(32) in type 1^WT-TN^ and type 2^WT-cold^, respectively. In the double-quinone (2Q) setups an additional CoQ_9_ was modeled at shallow sites 2/3 (see Table S 4). Note, we have used the Q site nomenclature described in in ref(32). The sites 1, 1F and 2/3 in correspond to the sites 1, 2 and 4, respectively, predicted earlier based on classical MD simulations(46, 83, 84).

The QM regions comprised CoQ species together with the surrounding protein residues (Table S 5). The CoQ_1_ ligand was included in its entirety in the QM region, whereas CoQ_9_ was pruned at the C_11_-C_12_ bond, comprising the identical number of atoms in the QM region to the CoQ_1_ species. This procedure was done to make two quinone types equal in terms of QM interactions which is substantial when analyzing spin density distributions. The QM/MM bond for protein sidechains was introduced between the Cα-Cβ atoms. Different redox and spin states were studied by altering the charge and the multiplicity of the QM region. All QM regions (shown in Table S 5) had charge 1 in their native state. The one-electron reduced case was simulated as a doublet (multiplicity=2), and the two-electron reduced case was considered either a closed-shell singlet (multiplicity=1) or an open-shell triplet (multiplicity=3).

The protein system was incorporated into a lipid bilayer containing POPC, POPE, and TLCL lipids (with a proportion 50%, 36%, and 14%, respectively), and immersed into a water-ionic box with the dimensions 105 Å x 130 Å x 150 Å, and concentration of ions 100 mM. The model system comprised 167,540 atoms in total. The setup was initially minimized for 1000 steps using the conjugate gradient algorithm with constraints on protein and quinone heavy atoms, followed by the equilibration in an NPT ensemble for 10 ns with the same constraints. After that, the QM/MM minimization was performed for 200 steps, followed by the production run for 1 ps in an NPT ensemble. The temperature was maintained at 310 K with Langevin thermostat(85), and the pressure control was implemented using Langevin Piston Nose-Hoover barostat(86, 87). Verlet method was used to integrate the MD trajectories with the time step of 1 fs.

QM/MM MD simulations were performed with NAMD(88) (version v2.14) and ORCA (version 5.0.3) software packages. The protonation states of the titratable residues were determined from pKa calculations with PROPKA software(89). The Verlet scheme(90) was used for the treatment of non-bonded interactions with the cutoff, switching, and pairlist distances of 12 Å, 10 Å, and 14 Å, respectively. Long-range electrostatic interactions were described by the Particle Mesh Ewald (PME) method(81). MM region was parametrized by CHARMM36 force field(91).

The QM interactions were considered in the framework of density functional theory (DFT) with hybrid B3LYP functional(92, 93), def2-SVP basis set(94), and the SCF energy tolerance of 10^-8^ au. The additive electrostatic embedding scheme(88) was used to couple the QM region to the MM environment. VMD visualization tool(82) was utilized for analyzing the trajectories and plotting spin densities and creating figures.

Spin and charge populations on CoQ species were derived with the Mulliken method for population analysis(95). Electron transfer rate constants (k_ET_) were estimated with Moser Dutton ruler(96):

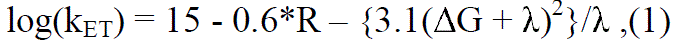

where k_ET_ is measured in s^-1^, R (edge-to-edge distance) is in Å, ΔG and λ (in eV) are free energy difference and reorganization energy, respectively. According to the previous studies of the electron transfer in proteins(*97*), they are set to: λ = 0.5 eV, ΔG = 0 eV.

